# Prevalence of pathogenic variants in DNA damage response genes in patients undergoing cancer risk assessment and reporting a personal history of early-onset renal cancer

**DOI:** 10.1101/2020.02.21.958256

**Authors:** Tiffiney R. Hartman, Elena V. Demidova, Randy W. Lesh, Lily Hoang, Marcy Richardson, Andrea Forman, Lisa Kessler, Virginia Speare, Erica A. Golemis, Michael J. Hall, Mary B. Daly, Sanjeevani Arora

## Abstract

**Purpose:** Pathogenic variants (PVs) in a number of genes are known to increase the risk of hereditary renal cancer (hRC). However, many early-onset RC (eoRC) patients undergoing genetic testing lack PVs in hRC genes; thus, their genetic risk remains undefined. To determine if PVs in DNA damage response (DDR) genes are enriched in a convenience sample of eoRC patients undergoing genetic testing.

**Materials and Methods:** Retrospective review of results for 844 unselected eoRC patients, undergoing genetic testing with a multi-gene cancer panel by Ambry Genetics [between July 2012 and December 2016]. The patients were tested with CancerNext and/or CancerNext Expanded panels for a variety of indications. Identified PVs were compared with patient characteristics.

**Results:** Mean age of RC diagnosis was 48 years [range 24-60]. In addition to eoRC, 57.9% patients tested reported at least one additional cancer; breast cancer being the most common (40.1% of females, 2.5% of males). PVs in cancer risk genes were identified in 12.8% of patients—3.7% in RC-specific genes, and 8.55% in DDR genes. DDR gene PVs were most commonly identified in *CHEK2, BRCA1/2*, and *ATM*. Among the 2.1% of patients with a *BRCA1/2* PV, <50% reported a personal history of hereditary breast/ovarian-associated cancer. No association between age of RC diagnosis, and prevalence of PVs in RC-specific or DDR genes was observed.

**Conclusions:** Multi-gene panel testing including DDR genes may provide a more comprehensive risk assessment in unselected eoRC patients, and their families. Validation in larger datasets is needed to characterize the association with eoRC.

## INTRODUCTION

Renal cancer (RC) often develops with no signs or symptoms and is referred to as the “silent disease” (1). While factors including smoking, environmental factors, obesity, and race have been linked to increased risk of RC, inherited factors are the most well-validated source of increased risk (1). Hereditary RC (hRC) syndromes, typically associated with early-onset disease and a clinically significant family history of cancer, result from germline pathogenic variants (PV) in high-penetrance ‘RC-specific’ genes including *VHL, MET, FLCN, TSC1, TSC2, FH, SDH, PTEN* and *BAP1* (2-4). Previous report of an early-onset RC (eoRC) cohort screened with an RC-specific panel found 6.1% of individuals had a PV in an RC-specific gene (4). However, for most eoRC patients a PV in an RC-specific gene is not identified, leaving many eoRC genetically undefined. Thus, there is a need to identify additional genes related to eoRC risk. Currently, there are no National Comprehensive Cancer Network (NCCN) guidelines for detection, prevention, or risk reduction in individuals who present with an eoRC but lack a PV in a defined RC-specific gene (5).

DNA damage response (DDR) genes play an important role in maintaining genome integrity, but also increase cancer risk (6). PVs in DDR genes confer susceptibility to a variety of cancer types, but are not typically considered risk factors for eoRC; however, in some cases, eoRCs arise in patients with personal or family histories of other forms of cancer. This population is particularly likely to be interested in evaluation of genetic risk. Further, germline PVs in some DDR genes have been observed in RC, including the DNA mismatch repair (Lynch syndrome) genes *MSH2* and *MLH1* in renal urothelial carcinoma, and *CHEK2* in advanced renal cell carcinoma (7-17). To address the hypothesis that PVs in DDR genes may contribute to the missing heritability of eoRC, we analyzed germline sequencing data from a cohort of 844 eoRC, many of whom had additional non-RC primary tumors or a family history of non-RC cancers.

## MATERIALS AND METHODS

### Ambry Genetics eoRC study cohort, and variant determination

Genomic DNA (blood or saliva) from 844 RC patients (≤60 years at diagnosis, specimens collected between July 2012-December 2016) was tested with Ambry Genetics (Konica Minolta, Aliso Viejo, California) germline panels (CancerNext versions 1-4, and CancerNext-Expanded versions 1 & 2, [**Table S1**]). Data was requested for all eoRC patients tested with multi-gene cancer panels, and patients were unselected for family history of cancer, personal history of cancers (apart from RC), multifocal tumors, or RC-subtype/stage. De-identified patient information was analyzed for genetic test results and personal and family medical histories. Classification of variants is based on ACMG recommendations for standards for interpretation and reporting of sequence variations, and variants are regularly deposited in ClinVar by Ambry Genetics. Variant classification was updated through March 31, 2018 for all data. Gene variants were designated as PV (see below for criteria), variant of uncertain significance (VUS), or negative/indeterminate. Given updating of test panels by Ambry, not all patients were tested for all genes.

The Fox Chase Cancer Center Institutional Review Board (IRB) approved all work (IRB-14-831). The patients in the study did not consent to sharing their raw DNA sequence data. Individuals were provided different versions of the panel over the course of the study (for information on the patient cohort, Ambry CancerNext and CancerNext Expanded panels, see Supplemental file and **Table S1**).

### Statistical Analysis

To identify potential correlations between PVs and characteristics such as tumor pathology, additional primary tumor type, and age of diagnosis, genes were combined into pathways/groups of interest, histology’s were grouped, and cancer types were grouped. Each individual was categorized as having a variant in one of the genes within the group or no variant in the group. Gene categories were used for comparison of RC diagnosis with a DDR gene or an RC-specific gene. For additional methods, see Supplemental file.

## RESULTS

### Patient characteristics

We first benchmarked the eoRC study cohort to the reported incidence of RC in SEER data for the general US population to provide context. In the study cohort, 40% of cases were between 50-59 years of age, and median age of diagnosis was 48 years. As expected, a higher percentage of RC cases were diagnosed between 20-44 years of age as compared to patients ≤60 diagnosed with RC in the general US population (SEER) (35%, versus 21.9%) (**Fig. 1A**). The study cohort was 67.1% female and 32.9% male (**Fig. 1B, Table 1**), versus 34.8% female and 65.2% male for the general US population prevalence of RC diagnosed ≤60 (**Fig. 1B**). Race/ethnicities in study cohort were 65.6% Caucasian, 5.8% African American/Black, 5.3% Ashkenazi Jewish, 7.6% Hispanic, 0.5% other, and 5.5% unknown (**Table 1**).

**Table 1.**
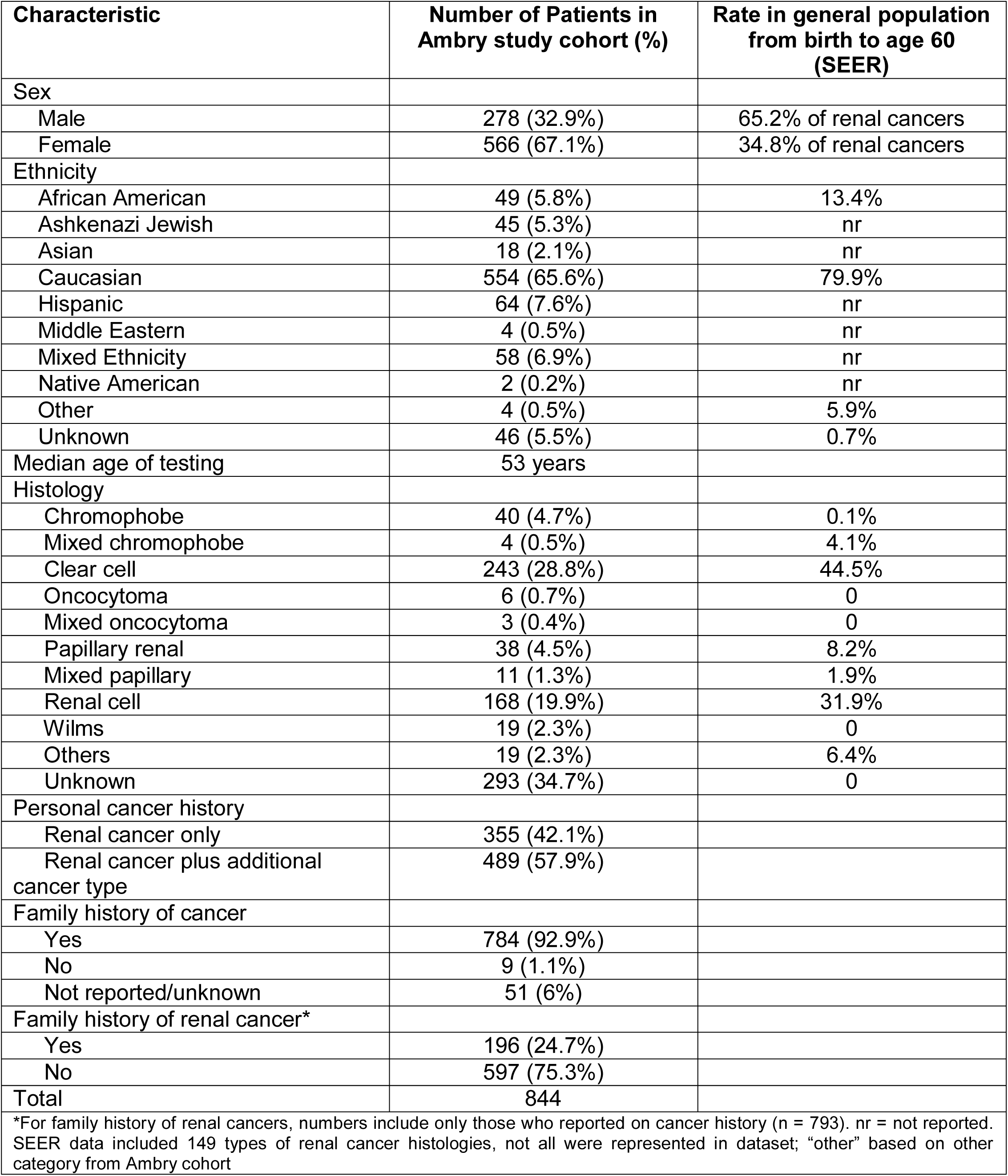
Demographics and clinical characteristics of RC patients in the Ambry Genetics study cohort. Demographics and clinical characteristics of the RC cases in the study cohort were compared to those of RC (from birth to age 60) in the SEER data. Personal and family history of cancer were reported for the cases in the study cohort.

**Figure 1.**
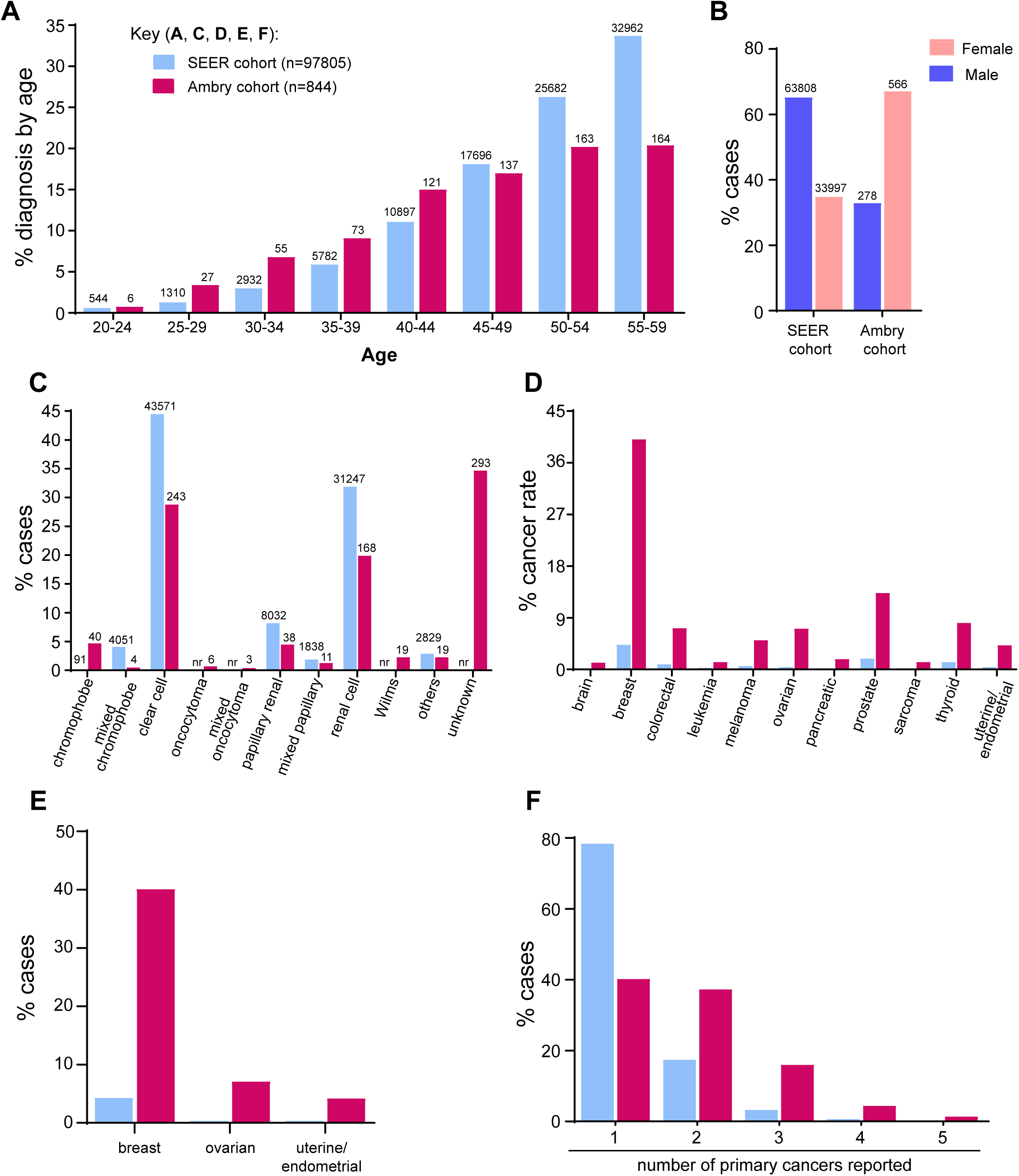
Patient characteristics. **A.** Age range of individuals diagnosed with RC ≤60 years in SEER cohort compared to the study cohort (n=746; of the remaining individuals in the study, 26 were diagnosed <19 years, 33 were diagnosed at 60 years, and 39 were excluded from the calculations as their age was reported as a wide range of years). **B.** Percentage of males and females diagnosed with RC ≤60 years in SEER compared to the study cohort (n = 844). **C.** The percentages of reported RC histology up to age 60 years in the SEER data (n=97,805) compared to the study cohort (n=844); not all histological subtypes reported in SEER were reported in the study cohort**. D.** The percentage of cancer incidence (at ≤60 years) in the general SEER population versus the study cohort. The SEER data reflect individuals reporting the indicated cancer type, not individuals with RC in addition to the indicated cancer type. **E.** The percentage of female specific cancers in SEER versus the study cohort. **F.** Percentage of different primary cancers reported (≤60 years) in SEER (n=97,795) versus the study cohort (n=844). Less than 0.4% not reported for figure clarity.

The tumor pathologies reported varied (**Fig. 1C and Table 1**). Clear cell constitutes 44.5% of all RCs in SEER, and was the most commonly reported histology in the eoRC cohort (243/844 = 28.8%). Renal cell (not defined, but likely to predominantly reflect clear cell) was also common (168/844 = 19.9%, **Fig. 1C and Table 1**). Papillary and chromophobe histology were each identified in ∼4-5% of the individuals (38/844 = 4.5% and 40/844 = 4.7%, respectively). Other histologies were identified rarely, but included Wilms tumor (19/844 = 2.3%) and oncocytoma (6/844 = 0.7%). For 34.7% of patients, the RC subtype was unknown.

### High incidence of other cancers in study cohort

57.9% (n=489/844) of the cases in the study cohort reported at least one additional primary cancer (**Fig. 1D, 1E, 1F, Table 1, Table S2**). Each of the primary cancer types is also represented at a higher level in the study cohort than in the general US population as reported by the SEER database (**Fig. 1D**). For female-specific cancers, 40.1% of females (227/566) also had breast cancer, in comparison to the 4.3% breast cancer rate in women ≤60 in the general population (SEER) (**Fig. 1D & 1E, Table S2**). The rate of additional primary cancer in the study cohort (57.9%) is much higher than the rate of each cancer type observed in SEER cases with eoRC (21.6%) (**Fig. 1F)**.

### Multi-gene cancer panel testing identifies PVs in DDR genes in the study cohort

The most common gene with PVs identified in the eoRC patients was the DDR gene *CHEK2* (19/844, 2.25%, **Fig. 2A, Table S3 & S4**), consistent with a recent report by Carlo *et al*.(13) Of patients with *CHEK2* PVs, 47.3% (n=9/19) had a common, highly damaging variant (c.1100delC, p.Thr367Met*fs*) that is associated with an increased risk for breast, prostate, colorectal and thyroid cancers (**Table S4**) (18-21).

**Figure 2.**
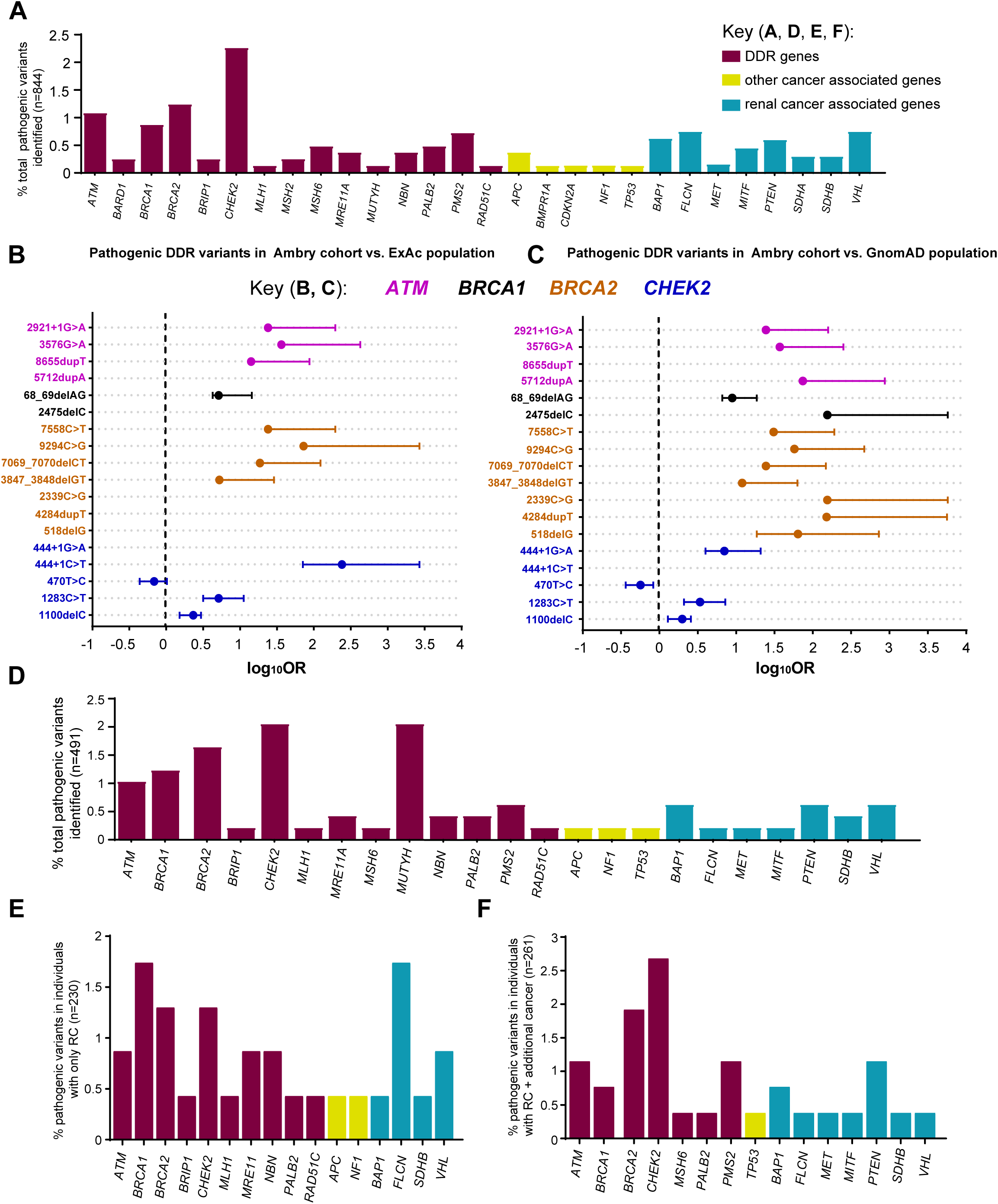
Enrichment of PVs in DDR genes. **A.** Cases with germline PVs in the entire cohort (n = 844). **A.** Red bars; DDR genes, yellow bars; other-cancer associated genes; blue bars; RC genes. *APC* variants identified in this study were all the moderate risk c.3920T>A, p.I1307K variant, and 5 of the 19 *CHEK2* variants were the moderate risk c.470T>C, p.I157T variant. *The individual with *MUTYH* was a compound heterozygote with two PVs. **B & C.** Odds of finding PVs in *ATM* (pink circle), *BRCA1* (black circle), *BRCA2* (orange circle), and *CHEK2* (blue circle) from study cohort versus control population, ExAc (**B)** and gnomAD (**C**). Data is presented as log10 odds ratio (OR), and log10 confidence intervals. Dotted black line; association with outcome i.e. OR>0 is enrichment in study cohort, OR=0 no difference in cohorts. PVs not found in gnomAD or ExAc are indicated by the absence of any data or PVs not listed from **Supplemental Table 4** were not found in the control population. **Note**: Computation of proportion or burden of individuals with all PVs in a specific gene(s) in the control population cannot be accurately performed as all PVs have not been defined, and while there is some agreement on which variants in a specific gene(s) are currently considered PVs, this is not true for all variants in that gene (as referenced in ClinVar, https://www.ncbi.nlm.nih.gov/clinvar/). **D.** Cases with germline PVs in the cohort tested for all genes in the study (n=491, 49 genes). **E.** Individuals with germline PVs who were diagnosed with only RC (n = 230/491). **F.** Individuals with germline PVs who were diagnosed with RC plus at least one additional primary cancer type (n=61/491). The color scheme as in **A**.

After *CHEK2*, PVs were most frequently observed in the DDR genes *BRCA2* (10/815, 1.23%), *ATM* (9/844, 1.07%) and *BRCA1* (7/815, 0.86%) (**Table S3**). We compared the overall frequency of PVs in *CHEK2, BRCA1/2*, and *ATM* to the control population in ExAc and gnomAD, representing individuals sequenced for disease-specific and population genetic studies (22, 23). Overall, PVs in each of these genes were more common in the study cohort versus the control populations (**Fig. 2B & 2C, Table S5A**). An outlier was the moderately damaging *CHEK2* c.470T>C p. I157T PV identified in 5 individuals in the study cohort, which was higher in the controls (gnomAD-OR, 0.60; 95% CI, 0.234-1.433; ExAc-OR, 0.72; 95% CI, 0.282-1.74). We compared the prevalence of all PVs in DDR genes presented in **Table S4**, from 844 cases, to controls from gnomAD.^23^ We found ∼4.8-fold enrichment of PVs in DDR genes in the study cohort versus the controls in gnomAD (8.4% vs. 1.8% respectively) (**Table S3 & S5B**; each DDR gene was corrected for number of patients assessed).

The overall gene variation rate in the full study cohort (n=844) is presented in **Table S1**. The full study cohort was not tested for all 49 genes. The largest panel was tested in 491 cases, and consisted of 49 genes, which included 15 RC-specific genes, and 34 other-cancer associated genes including 19 DDR genes (**Table S1**). Here, 12.8% (63/491) of cases had PVs. PVs were identified in one or more of the 15 genes not typically associated with RC in 9.16% cases (n=45/491, **Table S6**), versus 3.7% (n=18/491) with a PV in RC-specific genes (**Fig. 2D, Table S6**). Of the 15 genes not typically associated with RC, 12 were in DDR genes (8.55%, n=42/491 or 66.7%, n=42/63).

### DDR PVs are similarly enriched in patients diagnosed with eoRC alone, or with eoRC and other cancers

To test the hypothesis that DDR PVs might be associated with the phenotype of multiple primary cancers, we first analyzed PVs in eoRC cases with no additional primary cancer diagnosis. Among patients who only presented with eoRC (46.8%, n=230/491), PVs were identified in 13% of cases (n=30/230, **Fig. 2E**), which is approximately twice the reported frequency of PVs in RC-specific genes (4). Among this 13%, 8.7% (n=20/230) of PVs were in one of 10 DDR genes (*ATM, BRCA1/2, BRIP1, CHEK2, MLH1, MRE11A, NBN, PALB2, RAD51C*), 3.5% (n=8/230) were in one of 4 RC-specific genes (*BAP1, FLCN, SDHB, VHL*), and the remaining cases bore PVs in non-DDR genes associated with cancers other than RC (**Fig. 2E**).

We performed similar analysis for patients who presented with eoRC plus one or more additional cancer (53.2%, n=261/491). Among the patients with RC and at least one additional cancer, PVs were identified in 12.7% cases (33/261, **Fig. 2F).** Among these 12.7% of cases, PVs in other-cancer associated genes, including DDR genes, were found in 8.8% of cases (n=23/261), versus 3.8% (n=10/261) of cases with PVs in RC-specific genes. This population was also enriched for PVs in 7 DDR genes (8.4%, n=22/261, *ATM, BRCA1/2, CHEK2, MSH6, PALB2, PMS2*), versus PVs in 7 RC-specific genes (*BAP1, FLCN, MET, MITF, PTEN, SDHB, VHL*).

Overall, these data suggest that DDR PVs are enriched similarly in individuals diagnosed with eoRC alone or eoRC plus at least one additional primary cancer, but that the frequency of PVs in DDR genes, in either group, exceeded that in the control populations tested (gnomAD/ExAc) **(Fig. 2, Table S5A**). The specific PVs identified were similar in number and frequency to those identified in the full patient cohort (n=844), with *CHEK2* the most highly represented DDR gene (**Fig. 2**). To gain additional insight into the prevalence of these PVs in cancer patients, we surveyed ClinVar (https://www.ncbi.nlm.nih.gov/clinvar/), and found that multiple PVs from this study (**Table S4**) have been reported in hereditary cancer predisposing syndromes (HCPS, summarized in **Table S7**). HCPS reflects a pattern of cancers in a family characterized by earlier onset, with individuals not necessarily having the same tumor and/or having more than one primary tumor, and having tumors that are more likely to be multicentric.

### RC patients with BRCA PVs

Notably, 1.2% (10/815) of the eoRC cases had a *BRCA2* PV, and 0.9% (7/815) RC cases had a *BRCA1* PV (**Table 2, Table S3**). This included 1.7% (n = 6/355, **Table 2**) of the cases who presented with only eoRC. Interestingly, despite the fact that the cohort was 67.1% female, 47.1% (8/17) of the detected *BRCA1/2* PVs were in males (**Table 2**). Of the 17 RC cases with a *BRCA1 or BRCA2* PV, 8 (47.1%, 8/17) had an additional cancer associated with hereditary breast and ovarian cancer (HBOC) syndrome (breast, ovarian, prostate, pancreatic or melanoma), 3 had an additional non-HBOC cancer (17.6%, 3/17), and 6 presented with only eoRC (35.3%, 6/17) (**Table 2**). Family history was reported for 16 cases, and of those, 14/16 (87.5%) indicated that at least one family member had an HBOC-associated cancer. Of those with a *BRCA2* PV, 7/10 (70%) reported that at least one family member had RC (**Table 2**).

**Table 2.** Personal and family history of *BRCA1/2* positive patients in Ambry Genetics RC study cohort. Patient demographics, age of diagnosis, and cancer histories are listed. Family cancer histories listed for those who reported a family history.

| Gene | Gender | Age - RC | Personal cancer history | Age - HBOC syndrome cancer | Family history of cancer |
| --- | --- | --- | --- | --- | --- |
| *BRCA1* | M* | 51 | Renal | - | Breast, melanoma, gastric |
| *BRCA1* | F | 55 | Renal | - | Breast, ovarian |
| *BRCA1* | F | 45 | Renal | - | Breast, ovarian, gastric, lung |
| *BRCA1*<br>(+ *MSH6*) | M | 56 | Renal, prostate, CRC | 46 (p) | Ovarian, brain, uterine |
| *BRCA1* | F | 50 | Renal, ovarian, cervical | 63 | Not provided |
| *BRCA1* | F | 58 | Renal, breast, ovarian, sarcoma | 53 (b), 62 (o) | Throat, unknown |
| *BRCA1* | M | 0.5<br>(Wilm's tumor) | Renal, CNS, pheochromocytoma | - | Breast, prostate, gastric |
| *BRCA2* | F | 37 | Renal | - | Renal, breast, CRC |
| *BRCA2* | M | 52 | Renal | - | Renal, breast, pancreatic, CRC, thyroid, lung |
| *BRCA2* | M | 57 | Renal | - | No cancers |
| *BRCA2* | F | 51 | Renal, breast | 51 | Breast, pancreatic, thyroid, lung, skin |
| *BRCA2* | F | 60 | Renal, uterine | - | Renal, breast, pancreatic, prostate, sarcoma, bladder, thyroid, lymphoma |
| *BRCA2* | M | 46 | Renal, prostate | 53 | Breast, pancreatic, prostate |
| *BRCA2* | M | 54 | Renal, lung | - | Renal, breast, lung |
| *BRCA2* | F | 56 | Renal, breast | 59 | Renal, breast, ovarian, prostate, CRC, thyroid, lung |
| *BRCA2* | M | 56 | Renal, pancreatic | 56 (p) | Renal, breast, pancreatic, gastric |
| *BRCA2* | F | 51 | Renal, breast, CRC, uterine | 49 (b) | Renal, prostate, brain |

Abbreviations: b, breast cancer; CNS, central nervous system; CRC, colorectal cancer; HBOC syndrome, hereditary breast ovarian cancer syndrome; o, ovarian cancer; p, prostate cancer; RC, renal cancer; \*, Ashkenazi Jewish.

### No correlation between age of RC diagnosis and type of PV in eoRC

To determine if identification of specific classes of germline PV correlated with age of diagnosis in this cohort, genes were divided into four broad (overlapping) categories: all genes tested, RC-specific genes, non-RC associated genes (including DDR genes) and DDR genes (see Supplementary Methods). The groups were compared to median age at first RC diagnosis of <48 or ≥48 years. Given the invariable early-onset of Wilms tumor, the 20 individuals with this diagnosis were excluded from the analysis. Within this eoRC cohort, there was no significant association between age at diagnosis of RC and the type of PV for any of the four broad categories above (**Fig. 3A**).

**Figure 3.**
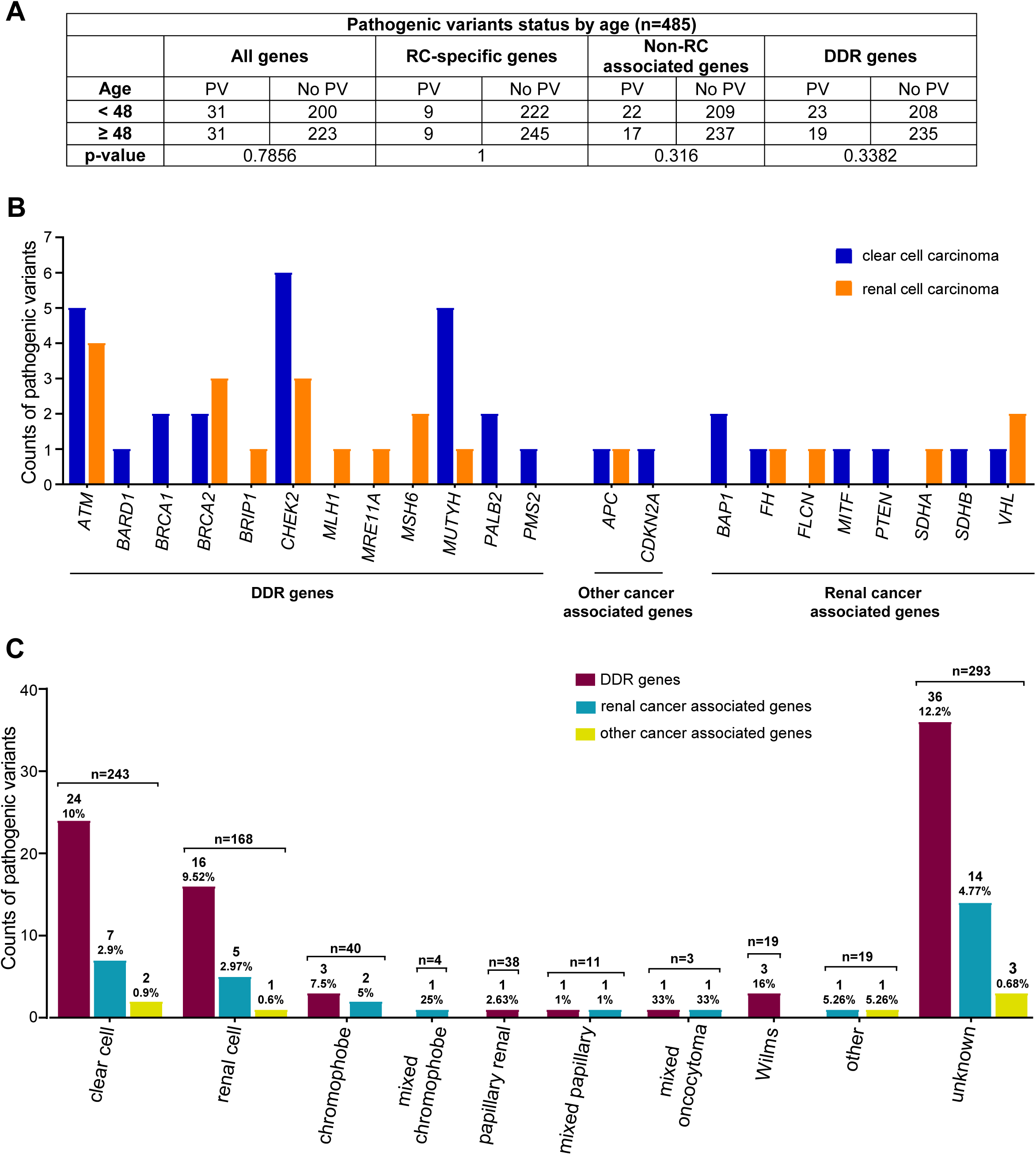
Identified PVs compared to age of RC diagnosis, and histology of RC. **A.** Statistical comparison of PVs in all genes, RC-specific genes, non-RC and DDR genes in younger individuals (<48) versus older individuals (≥48). N=485, these cases were tested for all 49 genes; all cases with Wilms tumor were removed. The results were non-significant (p>0.05) using two-sided Fisher’s exact tests. **B.** Counts of PVs by RC histology: clear cell (blue bar) and renal cell (orange bar) carcinoma in the study cohort. The’ renal cell’ subtype, is likely clear cell, but this cannot be confirmed. **C.** Counts of PVs by all RC histology observed in the study cohort. DDR genes (maroon bars), other-cancer associated genes (yellow bars), RC-specific genes (blue bars). The total number of individuals with a PV and percent PVs per gene category is shown above the bar. **B & C**. Includes counts from both homozygous and heterozygous carriers of *MUTYH*, and carriers of a *FH* variant that is currently considered to be pathogenic only in the compound heterozygous or homozygous state.

### Correlation of renal histologies with PVs in specific genes

Of the 243 clear cell cases in this cohort, 13.6% (33/243) had a PV, of which 2.9% were RC-associated PVs. Similar findings were observed for the cases described as renal cell carcinoma, 13.1% (22/168) had a PV, of which 2.4% were RC-associated. DDR gene PVs were found in 24/243 (∼10%) of clear cell cases, and in 16/168 (9.52%) of renal cell cases. **Fig. 3B & 3C** contrast the findings in clear cell and renal cell histology with the other non-clear cell histologies.

Odds ratios for identified PVs in specific renal tumor histologies were calculated. Specifically, for PVs in DDR genes, *ATM* correlated with mixed papillary histology, *BRCA1, MUTYH*, and *CHEK2* with Wilms tumor, *BRCA2* with papillary renal cell, and *CHEK2*, and *NBN* with chromophobe histology (**Table S8**). None of the associations were statistically significant.

## DISCUSSION

This study for the first time demonstrates that PVs in multiple DDR genes occur in patients with eoRC. Importantly, this study found that DDR PVs were highly represented both in cases diagnosed with eoRC and additional cancers, and also cases diagnosed with eoRC alone. Comparison with a large control population indicated that germline PVs in DDR genes were more common in this study cohort than in the control population, although further studies are required to confirm this finding and predict the penetrance of PVs in DDR genes for eoRC. We also found that germline testing using an RC-specific panel would have identified only 3.7% (18/491) of the RC cases with actionable PVs according to the NCCN recommended screening or management guidelines (NCCN Guidelines®), compared to the 9.16% (45/491) additional cases identified with the expanded panels.

Identification of germline DDR PVs can have specific implications for medical management and risks for the proband and the family. For example, 1.7% of cases diagnosed with eoRC alone had PVs in *BRCA1/2*, but not all of these cases had a family history strongly indicative of HBOC syndrome. Further, many of the specific PVs identified in this study have been annotated as relevant to HCPS, emphasizing the likely pathogenic nature. Our results support broader panel testing as a way to identify unexpected high-penetrant PVs in eoRC patients, when there is a personal or family history of additional cancers (especially an HBOC-syndrome cancer). This is important because identification of a *BRCA* PV can potentially change medical management; for instance, PARP inhibitor therapy is effective in tumors with *BRCA* PVs, including non-breast tumors (24, 25). Also, screening and prevention of HBOC-syndrome cancers will be increased significantly in the proband and family members found to have the same PV. In the study cohort, females were overrepresented, even though more males are typically diagnosed with RC (26). This difference may reflect likelihood of seeking genetic testing, given the frequency of HBOC in the cohort.

In RC, a number of germline PVs have been associated with treatment response. For example, bevacizumab with everolimus or erlotinib were added as treatment options for RC cases with germline PVs in *FH* (27, 28). Currently, the predictive and prognostic significance of PVs in DDR genes is not clinically defined for RC. There is an urgent need to study the biological impact of PVs in DDR genes in renal tissue. Such work may also lead to improved understanding of RC pathogenesis. Studies are in progress to assess cancer risk in different tissue types, and response to treatment due to a germline defect in DDR genes (29). A recent study showed that *VHL* inactivation in RC led to reduced expression of DDR genes (such as *BRCA1*/2), and thereby increased sensitivity to PARP inhibitors (30). These results indicate that RC tumors with DDR vulnerabilities may be responsive to PARP or other DDR inhibitors under development. Ongoing clinical trials are assessing the effect of PARP inhibitor, olaparib, in patients with somatic DDR variants in the setting of metastatic RC (<u>NCT03786796</u>). Finally, it is also important to evaluate the penetrance of DDR PVs to clinically define RC risk. These studies will assist in genetic counseling of RC patients and their families.

The limitations of this study include the following: This is a relatively small cohort, and all cases were not tested for all 49 genes. The cohort is a highly selected sample, as these patients likely had clinical characteristics (e.g. high rate of additional primary cancers) or family history that led to expanded panel testing. Although twice as many men are typically diagnosed with RC than women, the study cohort had more females than males. This may reflect the observation that women are more likely to pursue genetic testing than men, or the fact that 34.4% of cases also had a diagnosis of breast/ovarian/uterine cancer. Alternatively, men diagnosed with RC might be considered high-risk due to smoking or other environmental factors that lead their physician to be less suspicious of a hereditary component. A large percentage (34.7%) of tumors from the study cohort was listed as “unknown subtypes”, limiting comparison of PVs and RC histology types. Finally, matched (such as age, gender) comparisons were not possible using the control population, and we made no adjustment for population stratification. Differences in the study cohort, and control population ascertainment strategies and data collection (i.e. bioinformatic pipeline for variant calling/filtering, sequence coverage, race/ethnicities) prevent us from making any conclusions about the relationship between the PVs and RC risk. The comparisons performed in this manuscript were not adjusted for multiple testing.

## CONCLUSIONS

This study is the first to indicate a role for PVs in multiple DDR genes in eoRC. These results need to be validated in other large data sets. Additionally, to fully elucidate the biological relevance of DDR to RC, family and functional studies are needed as a next step to quantify the associated risks.

## DECLARATIONS

### Ethics approval and consent to participate

The Fox Chase Cancer Center Institutional Review Board (IRB) approved all work (IRB-14-831).

### Consent for publication

The patients in the study did not consent to sharing their raw DNA sequence data.

### Availability of data materials

All data generated or analysed during this study are included in this published article [and its supplementary files].

### Authors’ contributions

T.R.H., performed most studies, data analysis, writing; E.V.D., data analysis, figure preparation; V.S., L.H., sequence data, discussion; E.A.G., M.J.H., M.B.D., A.F., L.K., discussion & writing; S.A., study conception, discussion, planning & writing.

## Acknowldedgments

We express gratitude to Dr. Margie Clapper, for many helpful discussions and advice with this work. We thank members of the FCCC Risk Assessment Program and Department of Clinical Genetics, and Yan Zhou and Elizabeth Handorf in the FCCC Bioinformatics and Biostatistics Facility, for assisting with this work. Lisa Bealin (Risk Assessment Program, FCCC) for assisting with the IRB approval, and the FCCC Office of Research and Developmental Alliances for executing Data Use and License Agreements for work with Ambry Genetics.

