## Supplemental file for "Prevalence of pathogenic variants in DNA damage response genes in patients undergoing cancer risk assessment and reporting a personal history of early-onset renal cancer"

**Materials and Methods**

**Patient cohort: Ambry Genetics population**

Any de-identified personal or family history information including sex, ethnicity/race, age of cancer diagnosis, tumor histology, history of additional personal cancer, and history of family cancer and types was reported first as summarized data and later as de-identified individual case reports. For analysis comparing outcomes for RC-specific genes versus genes not typically associated with RC, we focused our statistical comparison on only those individuals who had CancerNext Expanded panel version 2 testing which analyzes all 49 genes including the RC-specific genes. There were 491 individuals who had the CancerNext Expanded version 2 test, and those individuals were used for this statistical comparison. For additional statistical test comparisons that analyzed the correlations between specific genes and categories such as tumor pathology or age, any individual who had been tested for that specific gene was included. Ambry Genetics follows strict criteria when categorizing variants as PV, Variant Likely Pathogenic (VLP), Variant Unknown Significance (VUS), Variant Likely Benign (VLB) and Benign (for details see https://www.ambrygen.com/clinician/our-scientific-excellence/variant-classification). All VUS were reviewed, but not reported in this study. All positive and negative data from germline testing of individuals diagnosed with renal cancer at age 60 or younger that was provided was utilized in our study.

**Genetic analysis with Ambry CancerNext and CancerNext Expanded panels**

Individuals were provided different versions of the panel over the course of the study (see **Table S1**). The number of genes in the panels ranged from the smallest CancerNext panel Version 1 which include 22 genes (*APC, ATM, BARD1, BRIP1, BMPR1A, CDH1, CHEK2, EPCAM, MLH1, MRE11A, MSH2,* *MSH6, MUTYH, NBN, PALB2, PMS2, PTEN, RAD50, RAD51C, SMAD4, STK11, TP53*) to the largest CancerNext Expanded Version 2 panel, which contained 49 genes (*APC, ATM, BAP1, BARD1, BRCA1, BRCA2, BRIP1, BMPR1A, CDH1, CDK4, CDKN2A, CHEK2, EPCAM, FH, FLCN, GREM1, MAX, MEN1, MET, MITF, MLH1, MRE11A, MSH2, MSH6, MUTYH, NBN, NF1, PALB2, PMS2, POLD1, POLE, PTEN, RAD50, RAD51C, RAD51D, RET, SDHA, SDHAF2, SDHB, SDHC, SDHD, SMAD4, SMARCA4, STK11, TMEM127, TP53, TSC1, TSC2, VHL*). The DDR genes identified in germline testing of this cohort are underlined (1).

For the CancerNext and CancerNext Expanded panels, genes (excluding *EPCAM* and *GREM1*) were analyzed by next generation or Sanger sequencing of all coding regions and at least 20 bases into flanking introns and 5’ and 3’ ends of the genes. Promoter regions were sequenced for *PTEN, MLH1,* and *MSH2*. Missense mutations outside of the exonuclease domains of *POLD1* and *POLE* were not reported. Additional details on specific testing methods are available at https://www.ambrygen.com/clinician/genetic-testing/28/oncology/cancernext-expanded.

**Control population in ExAc and gnomAD**

To compare the frequency of DDR PVs found in the study to that in the general population, our results were compared to the Exome Aggregation Consortium (ExAc) dataset of largely unrelated ~60,000 whole exome sequencing results, and to the Genome Aggregation database (gnomAD) dataset consisting of ~125,000 exomes and ~15,000 genomes (2, 3). These datasets are the most commonly used genomic data at the population-level.

**ClinVar Analysis**

ClinVar (<https://www.ncbi.nlm.nih.gov/clinvar/>), a database of medically relevant gene variants, was used to investigate all PVs in this study (retrieved on February 4, 2020). PVs that were not reported in ClinVar were noted as ‘*not reported’*. Associated conditions for each PV were categorized into hereditary cancer predisposing syndrome(s), condition(s) related to renal cancer, and any other condition(s). To further elucidate any PVs related to renal cancer, the search term “renal cancer” was queried, and the results were noted as “*associated with ClinVar search term ‘Renal Cancer.’”*

**Statistical Analysis**

We tested the hypothesis that different gene groups are associated with clinical characteristics of age at first kidney diagnosis of <48 years or ≥48 years, tumor histology, diagnosis of other cancer types, and whether kidney was the first primary cancer. To test the association between clinical parameters or membership of pathways and the presence of mutations, two-sided Fisher’s exact tests were used, and p-values ≤ 0.05 were considered significant. Odds ratios (OR) were calculated to determine the odds that an outcome had occurred given a particular variant, compared to the odds of the outcome occurring in the absence of that variant in the population tested. Finally, we queried the Surveillance, Epidemiology, and End Results-17 Registries database (SEER) database to find patients under 60 years old with kidney tumors and assess the distribution of their clinical characteristics (where available) (4).

Due to the evolving nature of the panels during the course of this study, each version included a different total number of genes, and analysis of each gene is based on the number of individuals whose test included that gene (**Table S1**). Only data from 491 individuals was considered for comparison of individuals with RC-specific genes compared to those with variants in genes not typically associated with RC, as the other individuals did not have all 49 genes analyzed. For statistical comparisons analyzing correlations between specific genes with various characteristics, all individuals who had been tested for that specific gene were included.

To identify potential correlations between PVs and characteristics such as tumor pathology, additional primary tumor type, and age of diagnosis, RC-specific genes, other cancer-associated genes, and DDR genes were combined into groups, and histologies were grouped. The categories for comparison of PVs and patient characteristics are as follows:

1. Known RC genes (*BAP1, FH, FLCN, MEN1, MET, MITF, PTEN, SDHA, SDHAF2, SDHB, SDHC, SDHD, TSC1, TSC2,* and *VHL* ) versus genes not typically associated with RC (*APC, ATM, BARD1, BRCA1, BRCA2, BRIP1, BMPR1A, CDH1, CDK4, CDKN2A, CHEK2, EPCAM, GREM1, MAX, MLH1, MRE11A, MSH2, MSH6, MUTYH, NBN, NF1, PALB2, PMS2, POLD1, POLE, RAD50, RAD51C, RAD51D, RET, SMAD4, SMARCA4, STK11, TEMEM127, TP53*) versus DDR genes alone *(ATM, BARD1, BRCA1, BRCA2, BRIP1, CHEK2, MLH1, MRE11A, MSH2, MSH6, MUTYH, NBN, PALB2, PMS2, POLD1, POLE, RAD50, RAD51C, RAD51D).*

2. Histology categories combined from the original categories: 1) Chromophobe, 2) Papillary renal, 3) Clear cell, 4) Wilms, 5) Renal cell (likely clear cell but cannot be confirmed), 6) Unknown, 7) Mixed papillary [clear cell/papillary, papillary/renal/chromophobe renal, papillary transitional, and sarcomatoid/papillary/clear cell], 8) Mixed chromophobe [chromophobe/oncocytoma, chromophobe/renal cell, clear cell/chromophobe, and clear cell/oncocytoma/chromophobe], 9) Oncocytoma, 10) Mixed oncocytoma [clear cell/oncocytoma, oncocytoma/collecting duct, and renal cell/oncocytoma], and 11) Others [included clear cell/sarcomatoid, collecting duct, mixed epithelial and stromal, mucinous tubular and spindle cell, multilocular cystic renal, neuroendocrine, renal cell/Wilms, renal cortical, sarcomatoid, transitional, urothelial and urothelial transitional].

**Results**

**Increased incidence of other cancers in RC Ambry cohort.**

In multiple cases, the individual reported more than one primary tumor in the same cancer type. For example, an individual with a reported 6 total primary tumors listed only RC and brain (hemangioblastoma), but had individually reported primary RCs that occurred at ages 37, 41, 43, and 44, 44 (bilateral), which accounted for all 6 primary cancers. This individual was found to have a pathogenic variant in *VHL* (**Table S4**).

**Multi-gene panel testing identifies PVs in DDR genes in the study cohort**

Among the less understood PVs in DDR genes, *MUTYH* PVs are of particular interest because cancer risk has been defined for homozygous carriers, but not for heterozygous carriers (5). We identified an additional 1.7% (n=14/844) of RC patients as heterozygous carriers of *MUTYH* PVs **(Table S3 & S4**). Two additional individuals were heterozygous for *MUTYH* but also found to have a PV in another gene (one with a *BRCA1* PV, the other with a *SDHB* PV). While PVs in *FH* confer an autosomal dominant risk for renal cancer, the two cases (0.3%, n=2/680) identified in this study were carriers of a specific gene variant, *FH* c.1431_1433dupAAA, p.K477DUP, which is currently considered to be pathogenic only in the compound heterozygous or homozygous state **(Table S3 & S4) (6).**

**Correlation of renal histologies with PVs in specific genes.**

Odds ratios for identified PVs in specific genes were either more (OR >1) or less (OR <1) likely than would be expected by chance alone to be found with specific histology types (**Table S8**). PVs in *ATM* and *SDHB* correlated with mixed oncocytoma histology. Additional strong ORs included *MITF* with mixed chromophobe histology, *ATM* with mixed papillary, *BRCA1, MUTYH,* and *CHEK2* with Wilms tumor, *BRCA2* with papillary renal cell, and *CHEK2, FLCN, NBN*, and *PTEN* with chromophobe (**Table S8**). These data reflect previous publications linking *SDHB* variants with renal oncocytomas and *BRCA1* mutations with Wilms tumors, and indicate these results may link variants in specific genes with renal cancer pathology subsets (7, 8).

**Supplemental Tables**

**Supplemental Table 1**. **Genes in each of the six multi-gene cancer panels included in this study**. Genes in each panel are listed. Panel versions are: CancerNext versions 1-4, CancerNext Expanded versions 1 and 2. The known RC genes are shown in blue, and other cancer genes in black. DDR genes are bolded in black.

**Supplemental Table 2. Personal history of additional primary cancers in the renal cancer study cohort.** Personal history of the Ambry cohort was reported and then compared to the general population (from birth to age 60, SEER data).

**Supplemental Table 3. Identified variants and classification in renal cancer patients in the study cohort: Full data set (n=844).** The results for all tested genes and the number of patients tested are reported. The classification for each gene finding is reported as a pathogenic variant (PV) or a variant of uncertain significance (VUS).

**Supplemental Table 4. Overall summary and list of all pathogenic variants (PV) reported with patient clinical characteristics.** Genes and pathogenic variants identified in those genes are summarized. Zygosity, pathogenicity classification of the variant, tumor histology, and personal cancer history are also listed for all of the patients in the Ambry cohort identified to have a PV.

**Supplemental Table 5. Comparison of germline pathogenic variants (PVs) in the control population. A.** Germline pathogenic variants in the top 4 genes (*CHEK2, BRCA1/2* and *ATM*) identified in the Ambry cohort were compared with two control populations (ExAc and gnomAD). The odds ratios and confidence intervals are reported. **B.** Pathogenic variants in all DDR genes in the study cohort (n=844) were compared with those detected in the control population in gnomAD. The percent of each pathogenic variant detected was calculated by using the counts observed versus the total individuals assessed in gnomAD. For this comparison, *MUTYH* heterozygotes were removed.

**Supplemental Table 6. Identified variants and classification in renal cancer patients in the study cohort tested for all 49 genes (n=491).** The results for all tested genes and the number of patients tested are reported. The classification for each gene finding is reported as a pathogenic variant.

**Supplemental Table 7. Survey of ClinVar for previous report on PVs identified in the study.** Conditions in which the PVs have been reported is summarized.

**Supplemental Table 8. Correlation of pathogenic variants and renal histology type.** Odds ratios are reported for Ambry cohort individuals that were diagnosed with renal cancer (n=844).

**Supplemental Table 1. Genes tested in each version of CancerNext and CancerNext Expanded panel tests between March 2013 and December 2016, classification updated through March 2018.**

| **CancerNext** | | | | **CancerNext Expanded** | |  |
| --- | --- | --- | --- | --- | --- | --- |
| **v1** | **v2** | **v3** | **v4** | **v1** | **v2** |  |
| *APC* | *APC* | *APC* | *APC* | *APC* | *APC* |  |
| ***ATM*** | ***ATM*** | ***ATM*** | ***ATM*** | ***ATM*** | ***ATM*** |  |
|  |  |  |  |  | *BAP1* |  |
| ***BARD1*** | ***BARD1*** | ***BARD1*** | ***BARD1*** | ***BARD1*** | ***BARD1*** |  |
|  | ***BRCA1*** | ***BRCA1*** | ***BRCA1*** | ***BRCA1*** | ***BRCA1*** |  |
|  | ***BRCA2*** | ***BRCA2*** | ***BRCA2*** | ***BRCA2*** | ***BRCA2*** |  |
| ***BRIP1*** | ***BRIP1*** | ***BRIP1*** | ***BRIP1*** | ***BRIP1*** | ***BRIP1*** |  |
| *BMPR1A* | *BMPR1A* | *BMPR1A* | *BMPR1A* | *BMPR1A* | *BMPR1A* |  |
| *CDH1* | *CDH1* | *CDH1* | *CDH1* | *CDH1* | *CDH1* |  |
|  |  | *CDK4* | *CDK4* | *CDK4* | *CDK4* |  |
|  |  | *CDKN2A* | *CDKN2A* | *CDKN2A* | *CDKN2A* |  |
| ***CHEK2*** | ***CHEK2*** | ***CHEK2*** | ***CHEK2*** | ***CHEK2*** | ***CHEK2*** |  |
| *EPCAM* | *EPCAM* | *EPCAM* | *EPCAM* | *EPCAM* | *EPCAM* |  |
|  |  |  |  | *FH* | *FH* |  |
|  |  |  |  | *FLCN* | *FLCN* |  |
|  |  |  | *GREM1* |  | *GREM1* |  |
|  |  |  |  | *MAX* | *MAX* |  |
|  |  |  |  |  | *MEN1* |  |
|  |  |  |  | *MET* | *MET* |  |
|  |  |  |  | *MITF* | *MITF* |  |
| ***MLH1*** | ***MLH1*** | ***MLH1*** | ***MLH1*** | ***MLH1*** | ***MLH1*** |  |
| ***MRE11A*** | ***MRE11A*** | ***MRE11A*** | ***MRE11A*** | ***MRE11A*** | ***MRE11A*** |  |
| ***MSH2*** | ***MSH2*** | ***MSH2*** | ***MSH2*** | ***MSH2*** | ***MSH2*** |  |
| ***MSH6*** | ***MSH6*** | ***MSH6*** | ***MSH6*** | ***MSH6*** | ***MSH6*** |  |
| ***MUTYH*** | ***MUTYH*** | ***MUTYH*** | ***MUTYH*** | ***MUTYH*** | ***MUTYH*** |  |
| ***NBN*** | ***NBN*** | ***NBN*** | ***NBN*** | ***NBN*** | ***NBN*** |  |
|  |  | *NF1* | *NF1* | *NF1* | *NF1* |  |
| ***PALB2*** | ***PALB2*** | ***PALB2*** | ***PALB2*** | ***PALB2*** | ***PALB2*** |  |
| ***PMS2*** | ***PMS2*** | ***PMS2*** | ***PMS2*** | ***PMS2*** | ***PMS2*** |  |
|  |  |  | ***POLD1*** |  | ***POLD1*** |  |
|  |  |  | ***POLE*** |  | ***POLE*** |  |
| *PTEN* | *PTEN* | *PTEN* | *PTEN* | *PTEN* | *PTEN* |  |
| ***RAD50*** | ***RAD50*** | ***RAD50*** | ***RAD50*** | ***RAD50*** | ***RAD50*** |  |
| ***RAD51C*** | ***RAD51C*** | ***RAD51C*** | ***RAD51C*** | ***RAD51C*** | ***RAD51C*** |  |
|  |  | ***RAD51D*** | ***RAD51D*** | ***RAD51D*** | ***RAD51D*** |  |
|  |  |  |  | *RET* | *RET* |  |
|  |  |  |  | *SDHA* | *SDHA* |  |
|  |  |  |  | *SDHAF2* | *SDHAF2* |  |
|  |  |  |  | *SDHB* | *SDHB* |  |
|  |  |  |  | *SDHC* | *SDHC* |  |
|  |  |  |  | *SDHD* | *SDHD* |  |
| *SMAD4* | *SMAD4* | *SMAD4* | *SMAD4* | *SMAD4* | *SMAD4* |  |
|  |  |  | *SMARCA4* |  | *SMARCA4* |  |
| *STK11* | *STK11* | *STK11* | *STK11* | *STK11* | *STK11* |  |
|  |  |  |  | *TMEM127* | *TMEM127* |  |
| *TP53* | *TP53* | *TP53* | *TP53* | *TP53* | *TP53* |  |
|  |  |  |  | *TSC1* | *TSC1* |  |
|  |  |  |  | *TSC2* | *TSC2* |  |
|  |  |  |  | *VHL* | *VHL* |  |
| **29** | **14** | **55** | **66** | **189** | **491** | **Total 844** |

*CancerNext versions 1-4, CancerNext Expanded versions 1 and 2. Blue font – RC-specific genes, black font- other cancer genes, and **bold black**- DDR genes.

**Supplemental Table 2. Personal history of primary cancers in the study cohort**

| **Cancer type** | **Number of patients** | **% of patients in Ambry Renal Cancer cohort^$^** | **Rate in general population (SEER) up to age 60 (%)** |
| --- | --- | --- | --- |
| renal | 844/844 | 100% | 0.5% |
| breast (female) | 227/566 | 40.1% | 4.3% |
| breast (male) | 7/278 | 2.5% | * |
| ovarian (female) | 40/566 | 7.1% | 0.4% |
| colorectal | 61/844 | 7.2% | 0.9% |
| uterine/endometrial (female) | 24/566 | 4.2% | 0.4% |
| pancreatic | 15/844 | 1.8% | 0.2% |
| thyroid | 68/844 | 8.1% | 1.3% |
| gastric | 3/844 | 0.4% | 0.1% |
| leukemia | 11/844 | 1.3% | 0.3% |
| brain | 10/844 | 1.2% | 0.2% |
| sarcoma | 11/844 | 1.3% | 0.1% |
| adrenal | 1/844 | 0.1% | - |
| small Intestine | 2/844 | 0.2% | 0.1% |
| prostate (male) | 37/278 | 13.3% | 1.9% |
| biliary tract | 2/844 | 0.2% | 0.2% |
| ureter | 0/844 | 0.0% | - |
| melanoma | 43/844 | 5.1% | 0.6% |
| CNS other | 2/844 | 0.2% | - |
| Other | 121/844 | 14.3% | - |
|  |  | **Total cases= 844** |  |
| Multiple patients listed more than 1 type of cancer primary (in addition to renal), so the total number of patients in each category adds up to a higher percentage than the 57.9% of individuals with renal cancer plus additional cancer(s). The rate of cancer in the general population is based off of the listed cancer type, not individuals with renal AND that additional cancer; *not calculated, very rare. Total number of males = 278. Total number of females = 566^. $^percentages for gender specific cancers calculated for just male or female individuals | | | |

**Supplemental Table 3. Identified variants and classification in patients in the study cohort: Full data set**

| **Mutation category/gene frequency** | | | | | |
| --- | --- | --- | --- | --- | --- |
| **Gene** | **PV** | **PV%** | **VUS** | **VUS %** | **Total Cases** |
| ***CHEK2*** | 19 | 2.25 | 27 | 3.20 | 844 |
| ***BRCA2*** | 10 | 1.23 | 16 | 1.96 | 815 |
| ***ATM*** | 9 | 1.07 | 42 | 4.86 | 844 |
| ***BRCA1*** | 7 | 0.86 | 5 | 0.61 | 815 |
| ***PALB2*** | 4 | 0.47 | 13 | 1.54 | 844 |
| ***MRE11A*** | 3 | 0.36 | 7 | 0.83 | 844 |
| ***NBN*** | 3 | 0.36 | 11 | 1.30 | 844 |
| ***BARD1*** | 2 | 0.24 | 9 | 1.07 | 844 |
| ***BRIP1*** | 2 | 0.24 | 10 | 1.07 | 844 |
| *CDKN2A* | 1 | 0.12 | 4 | 0.50 | 801 |
| *EPCAM* | 0 | 0.00 | 0 | 0.00 | 844 |
| *NF1* | 1 | 0.12 | 6 | 0.75 | 801 |
| *BMPR1A* | 1 | 0.12 | 7 | 0.83 | 844 |
| ***MLH1*** | 1 | 0.12 | 7 | 0.83 | 844 |
| ***MSH2*** | 2 | 0.24 | 16 | 1.90 | 844 |
| ***MSH6*** | 4 | 0.47 | 18 | 2.13 | 844 |
| ***MUTYH*** | 1  (16 carriers) | 0.12  (2.13) | 4 | 0.47 | 844 |
| ***RAD51C*** | 1 | 0.12 | 12 | 1.42 | 844 |
| *TP53* | 1 | 0.12 | 4 | 0.47 | 844 |
| *APC* | 3 | 0.36 | 15 | 1.78 | 844 |
| *CDH1* | 0 | 0.00 | 8 | 0.95 | 844 |
| *CDK4* | 0 | 0.00 | 2 | 0.25 | 801 |
| ***PMS2*** | 7 | 0.71 | 22 | 2.61 | 844 |
| ***POLD1*** | 0 | 0.00 | 4 | 0.72 | 556 |
| ***POLE*** | 0 | 0.00 | 9 | 1.62 | 556 |
| ***RAD50*** | 0 | 0.00 | 10 | 1.18 | 844 |
| ***RAD51D*** | 0 | 0.00 | 4 | 0.50 | 801 |
| *RET* | 0 | 0.00 | 12 | 1.76 | 680 |
| *SMAD4* | 0 | 0.00 | 8 | 0.95 | 844 |
| *SMARCA4* | 0 | 0.00 | 5 | 0.90 | 556 |
| *STK11* | 0 | 0.00 | 4 | 0.47 | 844 |
| *TMEM127* | 0 | 0.00 | 1 | 0.15 | 680 |
| *GREM1* | 0 | 0.00 | 0 | 0.00 | 556 |
| *MAX* | 0 | 0.00 | 0 | 0.00 | 680 |
| *FLCN* | 5 | 0.74 | 7 | 1.03 | 680 |
| *VHL* | 5 | 0.74 | 3 | 0.44 | 680 |
| *BAP1* | 3 | 0.61 | 5 | 1.02 | 491 |
| *PTEN* | 5 | 0.59 | 6 | 0.71 | 844 |
| *MITF* | 3 | 0.44 | 0 | 0.00 | 680 |
| *SDHA* | 2 | 0.29 | 8 | 1.18 | 680 |
| *SDHB* | 2 | 0.29 | 3 | 0.44 | 680 |
| *MET* | 1 | 0.15 | 10 | 1.47 | 680 |
| **Gene** | **PV** | **PV%** | **VUS** | **VUS %** | **Total Cases** |
| *FH* | 0  (2 carriers) | 0.00  (0.29) | 4 | 0.59 | 680 |
| *MEN1* | 0 | 0.00 | 2 | 0.41 | 491 |
| *SDHAF2* | 0 | 0.00 | 0 | 0.00 | 680 |
| *SDHC* | 0 | 0.00 | 2 | 0.29 | 680 |
| *SDHD* | 0 | 0.00 | 2 | 0.29 | 680 |
| *TSC1* | 0 | 0.00 | 5 | 0.74 | 680 |
| *TSC2* | 0 | 0.00 | 15 | 2.21 | 680 |
| Number and percentages of patients with pathogenic variants (PV) or variants of unknown significance (VUS). Blue font - RC-specific genes, black font- other cancer genes, and **bold black**- DDR genes. Total number of individuals tested for each gene is listed. | | | | | |

**Supplemental Table 4. Overall summary and list of all pathogenic variants reported with patient clinical characteristics**

| **ID #** | **^A^All genes** | **Gene 1** | **Variant** | **protein** | **Gene 2** | **Variant 2** | **Protein 2** | **^B^Zyg** | **^C^Primary** | **Primary** | **Pathogenicity** | **^D^Hist** |
| --- | --- | --- | --- | --- | --- | --- | --- | --- | --- | --- | --- | --- |
| 360278D | * | ***APC*** | c.3920T>A | p.I1307K |  |  |  | het |  |  | Moderate Risk | U |
| 427464E |  | ***APC*** | c.3920T>A | p.I1307K |  |  |  | het | colorectal |  | Moderate Risk | Clear cell |
| 499693E |  | ***APC*** | c.3920T>A | p.I1307K |  |  |  | het | breast | brain | Moderate Risk | Renal cell |
| 300401C |  | ***ATM*** | c.3712_3716DELTTATT | p.L1238KFS*6 | ***CHEK2*** | c.1100DELC | p.T367MFS*15 | Het/ het |  |  | Positive | Clear cell |
| 324223D |  | ***ATM*** | c.5712DUPA | p.S1905Ifs*25 |  |  |  | het |  |  | Positive | Clear cell |
| 403941E | * | ***ATM*** | c.2921+1G>A | NULL |  |  |  | het |  |  | Positive | Clear cell, papillary |
| 377115D | * | ***ATM*** | c.2260C>T | p.Q754* |  |  |  | het |  |  | Positive | Renal cell |
| 326417D |  | ***ATM*** | c.3402+2T>C | NULL |  |  |  | het |  |  | Positive | Clear cell |
| 418728E | * | ***ATM*** | c.8655DUPT | p.V2886CFS*10 |  |  |  | het | thyroid |  | Positive | Clear cell |
| 379068D | * | ***ATM*** | c.2839-3_2839DELTAGTINSGATACTA | NULL |  |  |  | het | thyroid |  | Positive | U |
| 262074C |  | ***ATM*** | c.3576G>A | p.K1192K |  |  |  | het | breast |  | Positive | Clear cell |
| 374611D | * | ***ATM*** | c.8147T>C | p.V2716A |  |  |  | het | colorectal | leukemia | Positive | Renal cell |
| 346077D | * | ***BAP1*** | c.1251-1G>A | NULL |  |  |  | het |  |  | Positive | U |
| 353167D | * | ***BAP1*** | c.1916_1917DELTG | p.V639GFS*3 |  |  |  | het | breast |  | Positive | U |
| 393589D | * | ***BAP1*** | c.1717DELC | p.L573WFS*3 |  |  |  | het | pancreatic | melanoma | Positive | Clear cell |
| 300345C |  | ***BARD1*** | EX9del | NULL |  |  |  | het | breast |  | Positive | U |
| 306209D |  | ***BARD1*** | c.539_540DELAT | p.Y180* |  |  |  | het | colorectal |  | Positive | Clear cell |
| 196114C |  | ***BMPR1A*** | c.420DELT | p.P140PFS*4 |  |  |  | het |  |  | Positive | U |
| 362726D | * | ***BRCA1*** | c.68_69DELAG | p.E23VFS*17 |  |  |  | het |  |  | Positive | Clear cell |
| 485314E | * | ***BRCA1*** | c.2475DELC | p.D825EFS*21 |  |  |  | het |  |  | Positive | Clear cell |
| 348785D | * | ***BRCA1*** | c.5207T>C | p.V1736A |  |  |  | het |  |  | Positive | U |
| 404309E |  | ***BRCA1*** | c.68_69DUPAG | p.E23EFS*9 | ***MSH6*** | EX5_6del | NULL | Het/ het | colorectal | prostate | Positive | U |
| 461082E | * | ***BRCA1*** | c.68_69DELAG | p.E23VFS*17 |  |  |  | het | ovarian |  | Positive | Clear cell |
| 492891E | * | ***BRCA1*** | c.4128_4129DELAA | p.S1377Rfs*3 |  |  |  | het | breast | ovarian; sarcoma | Positive | U |
| 345261D | * | ***BRCA1*** | c.68_69DELAG | p.E23VFS*17 | ***MUTYH*** | c.1187G>A | p.G396D | het | CNS | pheochromocytoma | Positive | Wilm's |
| 378758D | * | ***BRCA2*** | c.7558C>T | p.R2520* |  |  |  | het |  |  | Positive | Clear cell |
| 384946D | * | ***BRCA2*** | c.8575C>T | p.Q2859* |  |  |  | het |  |  | Positive | Papillary renal |
| 486321E | * | ***BRCA2*** | c.2731DELG | p.E911KFS*4 |  |  |  | het |  |  | Positive | Renal cell |
| 368979D | * | ***BRCA2*** | c.425G>A | p.S142N |  |  |  | het | breast |  | Positive | Renal cell |
| 373598D | * | ***BRCA2*** | c.2339C>G | p.S780* |  |  |  | het | uterine |  | Positive | Renal cell |
| 451319E | * | ***BRCA2*** | c.3847_3848DELGT | p.V1283KFS*2 |  |  |  | het | prostate |  | Positive | Renal cell |
| 326049D |  | ***BRCA2*** | c.7069_7070DELCT | p.L2357VFS*2 |  |  |  | het | lung |  | Positive | U |
| 458288E | * | ***BRCA2*** | c.518DELG | p.G173VFS*12 |  |  |  | het | breast |  | Positive | U |
| 407307E | * | ***BRCA2*** | c.4284DUPT | p.Q1429Sfs*9 |  |  |  | het | pancreatic |  | Positive | U |
| 188629B |  | ***BRCA2*** | c.9294C>G | p.Y3098* |  |  |  | het | breast | colorectal; uterine | Positive | U |
| 465722E | * | ***BRIP1*** | c.2108DELAINSTCC | p.K703IFS*3 |  |  |  | het |  |  | Positive | Renal cell |
| 275025C |  | ***BRIP1*** | c.2392C>T | p.R798* |  |  |  | het |  |  | Positive | U |
| 302273D |  | ***CDKN2A*** | c.202_203DELGCINSTT | NULL |  |  |  | het |  |  | Positive | Clear cell |
| 418115E | * | ***CHEK2*** | c.470T>C | p.I157T |  |  |  | het |  |  | Moderate Risk | Chromophobe |
| 405073E | * | ***CHEK2*** | c.470T>C | p.I157T |  |  |  | het | thyroid |  | Moderate Risk | U |
| 331880D |  | ***CHEK2*** | c.470T>C | p.I157T |  |  |  | het | breast |  | Moderate Risk | Clear cell |
| 344527D |  | ***CHEK2*** | c.470T>C | p.I157T |  |  |  | het | breast |  | Moderate Risk | Renal cell |
| 255664C |  | ***CHEK2*** | c.470T>C | p.I157T |  |  |  | carrier | breast |  | Moderate Risk | U |
| 301436D |  | ***CHEK2*** | c.1427C>T | p.T476M |  |  |  | het |  |  | Positive | Clear cell |
| 464152E | * | ***CHEK2*** | c.1100DELC | p.T367MFS*15 |  |  |  | het |  |  | Positive | Clear cell |
| 174995B |  | ***CHEK2*** | c.1100DELC | p.T367MFS*15 |  |  |  | het |  |  | Positive | Renal cell |
| 396237D | * | ***CHEK2*** | c.1283C>T | p.S428F |  |  |  | het |  |  | Positive | U |
| 370178D | * | ***CHEK2*** | c.1100DELC | p.T367MFS*15 |  |  |  | het | small intestine |  | Positive | Chromophobe |
| 383340D | * | ***CHEK2*** | c.1100DELC | p.T367MFS*15 |  |  |  | het | colorectal |  | Positive | Clear cell |
| 398153D | * | ***CHEK2*** | c.349A>G | p.R117G |  |  |  | het | ovarian |  | Positive | Clear cell |
| 338465D | * | ***CHEK2*** | c.1283C>T | p.S428F |  |  |  | het | breast |  | Positive | Renal cell |
| 274372C |  | ***CHEK2*** | c.1100DELC | p.T367MFS*15 |  |  |  | het | breast |  | Positive | U |
| 481514E |  | ***CHEK2*** | c.1100DELC | p.T367MFS*15 |  |  |  | het | breast |  | Positive | U |
| 354527D | * | ***CHEK2*** | c.1100DELC | p.T367MFS*15 |  |  |  | het | colorectal | sarcoma | Positive | U |
| 482994E | * | ***CHEK2*** | c.444+1G>A | NULL |  |  |  | het | blood | basal cell | Positive | U |
| 149663A |  | ***CHEK2*** | c.1100DELC | p.T367MFS*15 |  |  |  | het | ovarian | thyroid | Positive | Wilm's |
| 275831C |  | ***FH*** | c.1431_1433DUPAAA | p.K477DUP |  |  |  | carrier |  |  | FH Carrier | Clear cell |
| 355692D | * | ***FH*** | c.1431_1433DUPAAA | p.K477DUP |  |  |  | carrier |  |  | FH carrier | Renal cell |
| 487260E | * | ***FLCN*** | c.619-1G>A | NULL |  |  |  | het |  |  | Positive | Renal cell |
| 367152D | * | ***FLCN*** | c.1153DELC | p.Q385SFS*83 |  |  |  | het |  |  | Positive | U |
| 408980E | * | ***FLCN*** | c.1429C>T | p.R477* |  |  |  | het |  |  | Positive | U |
| 469865E | * | ***FLCN*** | c.619-1G>A | NULL |  |  |  | het |  |  | Positive | U |
| 359775D | * | ***FLCN*** | c.1285DUPC | p.H429PFS*27 |  |  |  | het | prostate |  | Positive | Chromophobe |
| 366979D | * | ***MET*** | c.3754T>C | p.Y1252H |  |  |  | het | bladder |  | Positive | Papillary transitional |
| 279471C |  | ***MITF*** | c.952G>A | p.E318K |  |  |  | het |  |  | Positive | Chromophobe, oncocytoma |
| 472581E | * | ***MITF*** | c.952G>A | p.E318K |  |  |  | het | uterine |  | Positive | Clear cell |
| 319704D |  | ***MITF*** | c.952G>A | p.E318K |  |  |  | het | thyroid |  | Positive | U |
| 376143D | * | ***MLH1*** | c.2252_2253DELAA | p.K751SFS*3 |  |  |  | het |  |  | Positive | Renal cell |
| 179334B |  | ***MRE11A*** | c.1090C>T | p.R364* |  |  |  | het |  |  | Positive | Renal cell |
| 366686D | * | ***MRE11A*** | c.1090C>T | p.R364* |  |  |  | het |  |  | Positive | U |
| 404175E | * | ***MRE11A*** | c.1090C>T | p.R364* |  |  |  | het |  |  | Positive | U |
| 278735C |  | ***MSH2*** | c.1216C>T | p.R406* |  |  |  | het | breast |  | Positive | Renal cell |
| 300405C |  | ***MSH2*** | EX1_7inv | NULL |  |  |  | het | colorectal |  | Positive | Urothelial |
| 277842C |  | ***MSH6*** | c.1238G>A | p.W413* |  |  |  | het |  |  | Positive | Clear cell |
| 369189D | * | ***MSH6*** | c.2731C>T | p.R911* |  |  |  | het | uterine |  | Positive | Renal cell |
| 168306B |  | ***MSH6*** | c.3261DUPC | p.F1088Lfs*5 |  |  |  | het | colorectal | prostate | Positive | Renal cell |
| 379092D | * | ***MUTYH*** | c.91DELG | p.A31PFS*27 |  |  |  | carrier |  |  | MUTYH Carrier | Clear cell |
| 388533D | * | ***MUTYH*** | c.536A>G | p.Y179C |  |  |  | carrier |  |  | MUTYH Carrier | U |
| 440163E |  | ***MUTYH*** | c.1187G>A | p.G396D |  |  |  | carrier |  |  | MUTYH Carrier | U |
| 185631B |  | ***MUTYH*** | c.536A>G | p.Y179C |  |  |  | carrier | colorectal |  | MUTYH Carrier | Clear cell |
| 468288E | * | ***MUTYH*** | c.1187G>A | p.G396D |  |  |  | carrier | pancreatic |  | MUTYH Carrier | Clear cell |
| 265151C |  | ***MUTYH*** | c.1187G>A | p.G396D |  |  |  | carrier | colorectal |  | MUTYH Carrier | U |
| 376921D | * | ***MUTYH*** | c.963DELG | p.S322AFS*86 |  |  |  | carrier |  |  | MUTYH Carrier | U |
| 382476D | * | ***MUTYH*** | c.536A>G | p.Y179C |  |  |  | carrier | breast |  | MUTYH Carrier | U |
| 390001D | * | ***MUTYH*** | c.358DELG | p.E120RFS*26 |  |  |  | carrier | breast |  | MUTYH Carrier | U |
| 391880D | * | ***MUTYH*** | c.1187G>A | p.G396D |  |  |  | carrier | breast |  | MUTYH Carrier | U |
| 326752D |  | ***MUTYH*** | c.1187G>A | p.G396D |  |  |  | carrier | breast | plasmacytoma | MUTYH Carrier | Renal cell |
| 332262D |  | ***MUTYH*** | c.1214C>T | p.P405L |  |  |  | carrier | breast | pancreatic | MUTYH Carrier | U |
| 443101E | * | ***MUTYH*** | c.462+2T>G | NULL |  |  |  | carrier | breast | ovarian | MUTYH Carrier | U |
| 350033D | * | ***MUTYH*** | c.1187G>A | p.G396D |  |  |  | carrier | colorectal | leukemia; sarcoma; lymphoma | MUTYH Carrier | U |
| 305063D |  | ***MUTYH*** | c.1187G>A | p.G396D | ***MUTYH*** | c.536A>G | p.Y179C | homo | colorectal |  | Positive | Clear cell |
| 389103D | * | ***NBN*** | c.2238DELC | p.Y746* |  |  |  | het |  |  | Positive | Chromophobe |
| 392099D | * | ***NBN*** | c.657_661DELACAAA | p.K219NFS*16 |  |  |  | het |  |  | Positive | U |
| 478293E |  | ***NBN*** | c.657_661DELACAAA | p.K219NFS*16 | ***PMS2*** | c.1738A>T | p.K580* | het/ het | breast |  | Positive | U |
| 389052D | * | ***NF1*** | 5'UTR_3'UTRdel | NULL |  |  |  | het |  |  | Positive | U |
| 442106E | * | ***PALB2*** | c.2711G>A | p.W904* |  |  |  | het |  |  | Positive | Clear cell |
| 366836D | * | ***PALB2*** | c.3113G>A | p.W1038* |  |  |  | het | breast |  | Positive | Clear cell |
| 325444D |  | ***PALB2*** | EX13_3'UTRdup | NULL |  |  |  | het | thyroid |  | Positive | U |
| 326260D |  | ***PALB2*** | c.758DUPT | p.S254Ifs*3 |  |  |  | het | breast |  | Positive | U |
| 403637E | * | ***PMS2*** | c.137G>T | p.S46I |  |  |  | het | sarcoma |  | Positive | Clear cell |
| 469026E | * | ***PMS2*** | c.736_741DELCCCCCTINSTGTGTGTGAAG | p.P246CFS*3 |  |  |  | het | melanoma |  | Positive | Renal cell |
| 383884D |  | ***PMS2*** | c.2155C>T | p.Q719* |  |  |  | het | uterine |  | Positive | U |
| 358014D |  | ***PMS2*** | c.137G>T | p.S46I | ***PMS2*** | c.137G>T | p.S46I | homo | colorectal | bladder | Positive | U |
| 437423E | * | ***PMS2*** | c.1A>G | p.M1? |  |  |  | het | prostate | throat | Positive | U |
| 371364D | * | ***PTEN*** | c.49C>T | p.Q17* |  |  |  | het | ovarian |  | Positive | Chromophobe |
| 303552D |  | ***PTEN*** | c.209+4_209+7DELAGTA | NULL |  |  |  | het | breast |  | Positive | Clear cell |
| 335970D | * | ***PTEN*** | c.48T>A | p.Y16* |  |  |  | het | breast |  | Positive | U |
| 456158E | * | ***PTEN*** | c.46DUPT | p.Y16Lfs*28 |  |  |  | het | thyroid |  | Positive | U |
| 193578B |  | ***PTEN*** | in1_3'UTRdel | NULL |  |  |  | het | lymphoma | testicular | Positive | U |
| 427742E | * | ***RAD51C*** | c.905-2_905-1DELAG | NULL |  |  |  | het |  |  | Positive | Clear cell |
| 316289D |  | ***SDHA*** | c.1432_1432+1DELGG | NULL |  |  |  | het | breast |  | Positive | U |
| 325023D |  | ***SDHA*** | c.91C>T | p.R31* |  |  |  | het | breast | colorectal | Positive | U |
| 362953D | * | ***SDHB*** | c.725G>A | p.R242H |  |  |  | het | pheochromocytoma |  | Positive | Clear cell |
| 335567D | * | ***SDHB*** | c.380T>G | p.I127S | ***MUTYH*** | c.536A>G | p.Y179C | het/ het |  |  | Positive | Oncocytoma, collecting duct |
| 362468D | * | ***TP53*** | c.827C>A | p.A276D |  |  |  | het | prostate | melanoma | Positive | Transitional |
| 301742D |  | ***VHL*** | 5'UTR_3'UTRdel | NULL |  |  |  | het |  |  | Positive | Renal cell |
| 330592D |  | ***VHL*** | c.266T>C | p.L89P |  |  |  | het |  |  | Positive | Clear cell |
| 423807E | * | ***VHL*** | c.238A>G | p.S80G |  |  |  | het |  |  | Positive | Renal cell |
| 496181E | * | ***VHL*** | c.232A>T | p.N78Y |  |  |  | het |  |  | Positive | Sarcomatoid |
| 400897E | * | ***VHL*** | c.481C>T | p.R161* |  |  |  | het | brain |  | Positive | U |

^A^Individuals with all 49 genes tested with CancerNext Expanded Version 2 are indicated with an asterisk.

^B^Zygosity: Het = heterozygous; Homo = homozygous; carrier = carrier of variant that is classified as nonpathogenic in the heterozygous state

^C^all individuals have a primary of renal cancer. Additional primaries are listed

^D^Histology, U = unknown

**Supplemental Table 5A. Comparison of pathogenic variants in *ATM*, *BRCA1/2* and *CHEK2* with control population. 5B. Comparison of pathogenic variants in all DDR genes in the study with control population.**

**5A**

| Study cohort vs. gnomAD |  |  |  |  |  |  |  |  |  |  |  |  |  |  |
| --- | --- | --- | --- | --- | --- | --- | --- | --- | --- | --- | --- | --- | --- | --- |
| Pathogenic variant (PV) | Study cohort - cases without the PV | Study cohort - cases with the PV | gnomAD - cases without the PV | gnomAD - cases with the PV | Odds Ratio | LCL | UCL | P-value | Log10 Odds Ratio | Log 10 LCL | Log 10 UCL | Ln Odds Ratio | Ln LCL | Ln UCL |
| NM_000051.3(ATM):c.3576G>A | 843 | 1 | 125559 | 4 | 37.2291 | 1.5340 | 286.0916 | 0.0329 | 1.5709 | 0.1858 | 2.4565 | 3.6171 | 0.4279 | 5.6563 |
| NM_000051.3(ATM):c.5712dupA | 843 | 1 | 125571 | 2 | 74.4568 | 2.5657 | 951.9962 | 0.0199 | 1.8719 | 0.4092 | 2.9786 | 4.3102 | 0.9422 | 6.8586 |
| NM_000051.3(ATM):c.2921+1G>A | 843 | 1 | 125646 | 6 | 24.8209 | 1.0949 | 185.0418 | 0.0458 | 1.3948 | 0.0394 | 2.2673 | 3.2117 | 0.0907 | 5.2206 |
| NM_007294.3(BRCA1):c.68_69delAG | 812 | 3 | 141163 | 58 | 8.9918 | 2.3810 | 27.7466 | 0.0053 | 0.9538 | 0.3768 | 1.4432 | 2.1963 | 0.8675 | 3.3231 |
| NM_007294.3(BRCA1):c.2475delC | 814 | 1 | 125455 | 1 | 154.2552 | 3.9990 | 5,932.6488 | 0.0129 | 2.1882 | 0.6020 | 3.7732 | 5.0386 | 1.3860 | 8.6882 |
| NM_000059.3(BRCA2):c.7558C>T | 814 | 1 | 125589 | 5 | 30.8575 | 1.3230 | 221.3283 | 0.0381 | 1.4894 | 0.1216 | 2.3450 | 3.4294 | 0.2799 | 5.3996 |
| NM_000059.3(BRCA2):c.2339C>G | 814 | 1 | 125435 | 1 | 154.2329 | 3.9983 | 5,931.7031 | 0.0129 | 2.1882 | 0.6019 | 3.7732 | 5.0385 | 1.3859 | 8.6881 |
| NM_000059.3(BRCA2):c.3847_3848delGT | 814 | 1 | 117907 | 12 | 12.0695 | 0.5720 | 75.4126 | 0.0857 | 1.0817 | (0.2426) | 1.8774 | 2.4907 | (0.5586) | 4.3230 |
| NM_000059.3(BRCA2):c.9294C>G | 814 | 1 | 141123 | 3 | 57.7515 | 2.2348 | 524.4119 | 0.0228 | 1.7616 | 0.3492 | 2.7197 | 4.0561 | 0.8042 | 6.2623 |
| NM_000059.3(BRCA2):c.7069_7070delCT | 814 | 1 | 141159 | 7 | 24.7877 | 1.1141 | 173.9529 | 0.0450 | 1.3942 | 0.0469 | 2.2404 | 3.2103 | 0.1080 | 5.1588 |
| NM_000059.3(BRCA2):c.4284dupT | 814 | 1 | 122212 | 1 | 150.5953 | 3.8956 | 5,779.2916 | 0.0132 | 2.1778 | 0.5906 | 3.7619 | 5.0146 | 1.3598 | 8.6620 |
| NM_000059.3(BRCA2):c.518delG | 814 | 1 | 15696 | 1 | 19.2805 | 0.5003 | 742.2899 | 0.0963 | 1.2851 | (0.3008) | 2.8706 | 2.9591 | (0.6925) | 6.6097 |
| NM_007194.3(CHEK2):c.1100delC | 835 | 9 | 139603 | 591 | 2.5460 | 1.2565 | 4.8760 | 0.0111 | 0.4059 | 0.0992 | 0.6881 | 0.9345 | 0.2283 | 1.5843 |
| NM_007194.3(CHEK2):c.470T>C | 839 | 5 | 139999 | 1391 | 0.5998 | 0.2354 | 1.4333 | 0.3763 | (0.2220) | (0.6282) | 0.1563 | (0.5112) | (1.4465) | 0.3600 |
| NM_007194.3(CHEK2):c.1283C>T | 842 | 2 | 140970 | 131 | 2.5560 | 0.4489 | 9.7547 | 0.1876 | 0.4076 | (0.3479) | 0.9892 | 0.9385 | (0.8010) | 2.2777 |
| NM_007194.3(CHEK2):c.444+1G>A | 843 | 1 | 141299 | 40 | 4.1903 | 0.2096 | 25.1233 | 0.2166 | 0.6222 | (0.6786) | 1.4001 | 1.4328 | (1.5626) | 3.2238 |
| Study cohort vs. ExAc |  |  |  |  |  |  |  |  |  |  |  |  |  |  |
| Pathogenic variant (PV) | Study cohort - cases without the PV | Study cohort - cases with the PV | ExAc - cases without the PV | ExAc - cases with the PV | Odds Ratio | LCL | UCL | P-value | Log10 Odds Ratio | Log 10 LCL | Log 10 UCL | Ln Odds Ratio | Ln LCL | Ln UCL |
| NM_000051.3(ATM):c.2921+1G>A | 843 | 1 | 60533 | 3 | 23.9458 | 0.9257 | 217.1996 | 0.0539 | 1.3792 | (0.0335) | 2.3369 | 3.1758 | (0.0772) | 5.3808 |
| NM_000051 (ATM):exon24:c.G3576A | 843 | 1 | 60562 | 2 | 35.9110 | 1.2374 | 459.1464 | 0.0407 | 1.5552 | 0.0925 | 2.6620 | 3.5810 | 0.2130 | 6.1294 |
| NM_000051.3 (ATM):c.8655dupT | 843 | 1 | 59979 | 5 | 14.2328 | 0.6101 | 102.0601 | 0.0804 | 1.1533 | (0.2146) | 2.0089 | 2.6555 | (0.4941) | 4.6256 |
| NM_007294.3(BRCA1):c.68_69delAG (Deletion) | 813 | 2 | 60457 | 29 | 23.8482 | 0.9232 | 216.6183 | 0.0540 | 0.7099 | (0.0604) | 1.2891 | 3.1717 | (0.0799) | 5.3781 |
| NM_000059:exon15 (BRCA2):c.C7558T:p.R2520X (LOF) | 843 | 1 | 60371 | 3 | 72.1986 | 1.8723 | 2,777.6896 | 0.0272 | 1.3775 | (0.0347) | 2.3357 | 4.2794 | 0.6272 | 7.9294 |
| NM_000059:exon25 (BRCA2):c.C9294G:p.Y3098X (LOF) | 814 | 1 | 58738 | 1 | 18.5909 | 0.7661 | 142.8787 | 0.0647 | 1.8585 | 0.2724 | 3.4437 | 2.9227 | (0.2664) | 4.9620 |
| NM_000059.3(BRCA2):c.7069_7070delCT (Deletion) | 814 | 1 | 60546 | 4 | 5.2523 | 0.2472 | 34.0965 | 0.1863 | 1.2693 | (0.1157) | 2.1550 | 1.6587 | (1.3976) | 3.5292 |
| NM_000059.3(BRCA2):c.3847_3848delGT | 814 | 1 | 47040 | 11 | 5.1274 | 0.8702 | 19.4563 | 0.0637 | 0.7203 | (0.6070) | 1.5327 | 1.6346 | (0.1390) | 2.9682 |
| NM_145862:exon3 (CHEK2):c.444+1C>T | 843 | 1 | 60682 | 1 | 72.0328 | 1.8678 | 2,770.9026 | 0.0272 | 1.8575 | 0.2713 | 3.4426 | 4.2771 | 0.6248 | 7.9269 |
| NM_007194.3(CHEK2):c.470T>C | 839 | 5 | 60175 | 497 | 0.7216 | 0.2825 | 1.7391 | 0.6969 | (0.1417) | (0.5490) | 0.2403 | (0.3263) | (1.2641) | 0.5534 |
| NM_007194.3(CHEK2):c.1283C>T | 842 | 2 | 59263 | 37 | 3.8045 | 0.6517 | 15.0521 | 0.1038 | 0.5803 | (0.1860) | 1.1776 | 1.3362 | (0.4282) | 2.7115 |
| NM_007194.3(CHEK2):c.1100delC | 835 | 9 | 58930 | 215 | 2.9542 | 1.4513 | 5.7897 | 0.0048 | 0.4704 | 0.1618 | 0.7627 | 1.0832 | 0.3725 | 1.7561 |

**5B**

| **Gene** | **Pathogenic variant (PV)** | **Allele count observed in gnomAD** | **Total individuals assessed in gnomAD** | **Percent detected in gnomAD** |
| --- | --- | --- | --- | --- |
| *ATM* | c.3712_3716DELTTATT | 0 | 125,573 | 0.00% |
| *ATM* | c.5712DUPA | 2 | 125,571 | 0.16% |
| *ATM* | c.2921+1G>A | 6 | 125,646 | 0.48% |
| *ATM* | c.2260C>T | 0 | 125,573 | 0.00% |
| *ATM* | c.3402+2T>C | 0 | 125,573 | 0.00% |
| *ATM* | c.8655DUPT | 0 | 125,573 | 0.00% |
| *ATM* | c.2839-3_2839DELTAGTINSGATACTA | 0 | 125,573 | 0.00% |
| *ATM* | c.3576G>A | 4 | 125,559 | 0.32% |
| *ATM* | c.8147T>C | 0 | 125,573 | 0.00% |
| *BRCA1* | c.68_69DELAG | 58 | 141,163 | 4.11% |
| *BRCA1* | c.2475DELC | 1 | 125,455 | 0.08% |
| *BRCA1* | c.5207T>C | 0 | 125,573 | 0.00% |
| *BRCA1* | c.68_69DUPAG | 0 | 125,573 | 0.00% |
| *BRCA1* | c.68_69DELAG | 0 | 125,573 | 0.00% |
| *BRCA1* | c.4128_4129DELAA | 0 | 125,573 | 0.00% |
| *BRCA1* | c.68_69DELAG | 0 | 125,573 | 0.00% |
| *BRCA2* | c.7558C>T | 5 | 125,589 | 0.40% |
| *BRCA2* | c.8575C>T | 0 | 125,573 | 0.00% |
| *BRCA2* | c.2731DELG | 0 | 125,573 | 0.00% |
| *BRCA2* | c.425G>A | 0 | 125,573 | 0.00% |
| *BRCA2* | c.2339C>G | 1 | 125,435 | 0.08% |
| *BRCA2* | c.3847_3848DELGT | 12 | 117,907 | 1.02% |
| *BRCA2* | c.7069_7070DELCT | 7 | 141,159 | 0.50% |
| *BRCA2* | c.518DELG | 1 | 15,696 | 0.64% |
| *BRCA2* | c.4284DUPT | 1 | 122,212 | 0.08% |
| *BRCA2* | c.9294C>G | 3 | 141,123 | 0.21% |
| *CHEK2* | c.470T>C | 1391 | 139,999 | 99.36% |
| *CHEK2* | c.1427C>T | 0 | 125,573 | 0.00% |
| *CHEK2* | c.1100DELC | 591 | 139,603 | 42.33% |
| *CHEK2* | c.1283C>T | 0 | 125,573 | 0.00% |
| *CHEK2* | c.349A>G | 0 | 125,573 | 0.00% |
| *CHEK2* | c.1283C>T | 131 | 140,970 | 9.29% |
| *CHEK2* | c.444+1G>A | 40 | 141,299 | 2.83% |
| *BARD1* | EX9del | 0 | 125,573 | 0.00% |
| *BARD1* | c.539_540DELAT | 3 | 141,018 | 0.21% |
| *BRIP1* | c.2108DELAINSTCC | 0 | 125,573 | 0.00% |
| *BRIP1* | c.2392C>T | 44 | 139,160 | 3.16% |
| *MLH1* | c.2252_2253DELAA | 0 | 125,573 | 0.00% |
| *MRE11A* | c.1090C>T | 0 | 125,573 | 0.00% |
| *MSH2* | c.1216C>T | 0 | 125,573 | 0.00% |
| *MSH2* | EX1_7inv | 0 | 125,573 | 0.00% |
| *MSH6* | EX5_6del | 0 | 125,573 | 0.00% |
| *MSH6* | c.1238G>A | 0 | 125,573 | 0.00% |
| *MSH6* | c.2731C>T | 1 | 15,699 | 0.64% |
| *MSH6* | c.3261DUPC | 1 | 15,563 | 0.64% |
| *MUTYH* | c.1187G>A | 3 | 140,573 | 0.21% |
| *NBN* | c.2238DELC | 0 | 125,573 | 0.00% |
| *NBN* | c.657_661DELACAAA | 57 | 141,066 | 4.04% |
| *PALB2* | c.2711G>A | 0 | 125,573 | 0.00% |
| *PALB2* | c.3113G>A | 17 | 141,378 | 1.20% |
| *PALB2* | EX13_3'UTRdup | 0 | 125,573 | 0.00% |
| *PALB2* | c.758DUPT | 6 | 125,683 | 0.48% |
| *PMS2* | c.137G>T | 48 | 138,590 | 3.46% |
| *PMS2* | c.736_741DELCCCCCTINSTGTGTGTGAAG | 0 | 125,573 | 0.00% |
| *PMS2* | c.2155C>T | 2 | 118,290 | 0.17% |
| *PMS2* | c.1A>G | 8 | 140,812 | 0.57% |
| *PMS2* | c.1738A>T | 0 | 125,573 | 0.00% |
| *RAD51C* | c.905-2_905-1DELAG | 3 | 125,664 | 0.24% |

**Supplemental Table 6. Identified variants and classification in renal cancer patients in the study cohort tested for all 49 genes (n=491).**

| **Mutation category/gene frequency** | | |
| --- | --- | --- |
| **Gene** | **PV** | **PV%** |
| ***CHEK2*** | 10 | 2.04 |
| ***BRCA2*** | 8 | 1.63 |
| ***BRCA1*** | 6 | 1.22 |
| ***ATM*** | 5 | 1.02 |
| ***PMS2*** | 3 | 0.61 |
| ***PALB2*** | 2 | 0.41 |
| ***MRE11A*** | 2 | 0.41 |
| ***NBN*** | 2 | 0.41 |
| ***BRIP1*** | 1 | 0.20 |
| *NF1* | 1 | 0.20 |
| ***MLH1*** | 1 | 0.20 |
| ***MSH6*** | 1 | 0.20 |
| ***MUTYH*** | 10 carriers | 2.04 |
| ***RAD51C*** | 1 | 0.20 |
| *TP53* | 1 | 0.20 |
| *APC* | 1 | 0.20 |
| *FLCN* | 5 | 1.02 |
| *VHL* | 3 | 0.61 |
| *BAP1* | 3 | 0.61 |
| *PTEN* | 3 | 0.61 |
| *SDHB* | 2 | 0.41 |
| *MITF* | 1 | 0.20 |
| *MET* | 1 | 0.20 |
| *FH* | 1 carrier | 0.20 |
| Number and percentages of patients with pathogenic variants (PV) in the 491 individuals tested with CancerNext Expanded version 2 including all 49 genes. Blue font - RC-specific genes, black font- other cancer genes, and **bold black**- DDR genes. Total number of individuals tested for each gene is listed. | | |

**Supplemental Table 7. Survey of ClinVar for previous report on PVs identified in the study.**

| **Gene** | **Variant** | **Hereditary Cancer Predisposing Syndrome(s)** | **Conditions(s) Related to Renal Cancer** | **Other Condition(s)** |
| --- | --- | --- | --- | --- |
| ***APC*** | c.3920T>A | Familial multiple polyposis syndrome, Familial adenomatous polyposis 1, Carcinoma of colon, Colorectal cancer, susceptibility to, Breast cancer, susceptibility to, Adenomatous polyposis coli, susceptibility to, Familial adenomatous polyposis, Hereditary cancer-predisposing syndrome |  |  |
| ***APC*** | c.3920T>A | Familial multiple polyposis syndrome, Familial adenomatous polyposis 1, Carcinoma of colon, Colorectal cancer, susceptibility to, Breast cancer, susceptibility to, Adenomatous polyposis coli, susceptibility to, Familial adenomatous polyposis, Hereditary cancer-predisposing syndrome |  |  |
| ***APC*** | c.3920T>A | Familial multiple polyposis syndrome, Familial adenomatous polyposis 1, Carcinoma of colon, Colorectal cancer, susceptibility to, Breast cancer, susceptibility to, Adenomatous polyposis coli, susceptibility to, Familial adenomatous polyposis, Hereditary cancer-predisposing syndrome |  |  |
| ***ATM*** | c.3712_3716DELTTATT | Hereditary cancer-predisposing syndrome, Ataxia-telangiectasia syndrome |  |  |
| ***ATM*** | c.5712DUPA | Hereditary cancer-predisposing syndrome, Ataxia-telangiectasia syndrome |  |  |
| ***ATM*** | c.2921+1G>A | Ataxia-telangiectasia syndrome, Hereditary cancer-predisposing syndrome, Ataxia-telangiectasia syndrome, Familial cancer of breast |  |  |
| ***ATM*** | c.2260C>T |  |  | not reported |
| ***ATM*** | c.3402+2T>C | Hereditary cancer-predisposing syndrome |  |  |
| ***ATM*** | c.8655DUPT |  |  | not reported |
| ***ATM*** | c.2839-3_2839DELTAGTINSGATACTA | Hereditary cancer-predisposing syndrome, Ataxia-telangiectasia syndrome |  |  |
| ***ATM*** | c.3576G>A | Hereditary cancer-predisposing syndrome, Ataxia-telangiectasia syndrome |  |  |
| ***ATM*** | c.8147T>C | Hereditary cancer-predisposing syndrome, Familial cancer of breast, Ataxia-telangiectasia syndrome |  |  |
| ***BAP1*** | c.1251-1G>A |  | Tumor susceptibility linked to germline BAP1 mutations |  |
| ***BAP1*** | c.1916_1917DELTG |  |  | not reported |
| ***BAP1*** | c.1717DELC |  |  | not reported |
| ***BARD1*** | EX9del |  |  | not reported |
| ***BARD1*** | c.539_540DELAT |  |  | not reported |
| ***BMPR1A*** | c.420DELT | Hereditary cancer-predisposing syndrome |  |  |
| ***BRCA1*** | c.68_69DELAG |  |  | not reported |
| ***BRCA1*** | c.2475DELC | Ovarian Neoplasms, Hereditary cancer-predisposing syndrome, Hereditary breast and ovarian cancer syndrome, Breast-ovarian cancer, familial 1 |  |  |
| ***BRCA1*** | c.5207T>C |  |  | not reported |
| ***BRCA1*** | c.68_69DUPAG |  |  | not reported |
| ***BRCA1*** | c.68_69DELAG |  |  | not reported |
| ***BRCA1*** | c.4128_4129DELAA | Hereditary cancer-predisposing syndrome, Hereditary breast and ovarian cancer syndrome, Breast-ovarian cancer, familial 1 |  |  |
| ***BRCA1*** | c.68_69DELAG |  |  | not reported |
| ***BRCA2*** | c.7558C>T | Hereditary breast and ovarian cancer syndrome, Hereditary cancer-predisposing syndrome, Neoplasm of the breast, Breast-ovarian cancer, familial 2, Breast-ovarian cancer, familial 1 |  |  |
| ***BRCA2*** | c.8575C>T | Hereditary cancer-predisposing syndrome, Hereditary breast and ovarian cancer syndrome, Breast-ovarian cancer, familial 2 |  |  |
| ***BRCA2*** | c.2731DELG | Breast-ovarian cancer, familial 2, Hereditary cancer-predisposing syndrome, Hereditary breast and ovarian cancer syndrome |  |  |
| ***BRCA2*** | c.425G>A | Hereditary cancer-predisposing syndrome, Breast-ovarian cancer, familial 2, Hereditary breast and ovarian cancer syndrome |  |  |
| ***BRCA2*** | c.2339C>G | Hereditary cancer-predisposing syndrome, Hereditary breast and ovarian cancer syndrome, Breast-ovarian cancer, familial 2 |  |  |
| ***BRCA2*** | c.3847_3848DELGT | Ovarian Neoplasms, Hereditary cancer-predisposing syndrome, Breast-ovarian cancer, familial 2, Breast and/or ovarian cancer, Hereditary breast and ovarian cancer syndrome, Familial cancer of breast |  |  |
| ***BRCA2*** | c.7069_7070DELCT | Breast-ovarian cancer, familial 2, Hereditary cancer-predisposing syndrome, Hereditary breast and ovarian cancer syndrome |  |  |
| ***BRCA2*** | c.518DELG | Breast-ovarian cancer, familial 2, Hereditary cancer-predisposing syndrome, Hereditary breast and ovarian cancer syndrome |  |  |
| ***BRCA2*** | c.4284DUPT | Breast-ovarian cancer, familial 2, Hereditary cancer-predisposing syndrome, Hereditary breast and ovarian cancer syndrome |  |  |
| ***BRCA2*** | c.9294C>G | Breast-ovarian cancer, familial 2, Breast-ovarian cancer, familial 1, Familial cancer of breast, Glioma susceptibility 3, Breast-ovarian cancer, familial 2, Pancreatic cancer 2, Malignant tumor of prostate, Medulloblastoma, Familial cancer of breast, Hereditary cancer-predisposing syndrome, Hereditary breast and ovarian cancer syndrome, Fanconi anemia, complementation group D1 | Wilms tumor 1 |  |
| ***BRIP1*** | c.2108DELAINSTCC | Hereditary cancer-predisposing syndrome, Familial cancer of breast, Fanconi anemia, complementation group J |  |  |
| ***BRIP1*** | c.2392C>T | Familial cancer of breast, Hereditary cancer-predisposing syndrome, Neoplasm of the breast, Neoplasm of ovary, Breast cancer, early-onset, Fanconi anemia, complementation group J |  | Tracheoesophageal fistula, BRIP1-Related Disorders |
| ***CDKN2A*** | c.202_203DELGCINSTT | Hereditary cutaneous melanoma, Hereditary cancer-predisposing syndrome |  |  |
| ***CHEK2*** | c.470T>C | Adrenocortical carcinoma, Gastrointestinal carcinoma, Hereditary cancer-predisposing syndrome, Prostate cancer, susceptibility to, Familial cancer of breast, Breast cancer, susceptibility to, Colon cancer, susceptibility to, Breast cancer, susceptibility to, Colorectal cancer, susceptibility to, Li-Fraumeni syndrome 2, Breast and colorectal cancer | Cancer of multiple types (associated with ClinVar search term "Renal Cancer") |  |
| ***CHEK2*** | c.470T>C | Adrenocortical carcinoma, Gastrointestinal carcinoma, Hereditary cancer-predisposing syndrome, Prostate cancer, susceptibility to, Familial cancer of breast, Breast cancer, susceptibility to, Colon cancer, susceptibility to, Breast cancer, susceptibility to, Colorectal cancer, susceptibility to, Li-Fraumeni syndrome 2, Breast and colorectal cancer | Cancer of multiple types (associated with ClinVar search term "Renal Cancer") |  |
| ***CHEK2*** | c.470T>C | Adrenocortical carcinoma, Gastrointestinal carcinoma, Hereditary cancer-predisposing syndrome, Prostate cancer, susceptibility to, Familial cancer of breast, Breast cancer, susceptibility to, Colon cancer, susceptibility to, Breast cancer, susceptibility to, Colorectal cancer, susceptibility to, Li-Fraumeni syndrome 2, Breast and colorectal cancer | Cancer of multiple types (associated with ClinVar search term "Renal Cancer") |  |
| ***CHEK2*** | c.470T>C | Adrenocortical carcinoma, Gastrointestinal carcinoma, Hereditary cancer-predisposing syndrome, Prostate cancer, susceptibility to, Familial cancer of breast, Breast cancer, susceptibility to, Colon cancer, susceptibility to, Breast cancer, susceptibility to, Colorectal cancer, susceptibility to, Li-Fraumeni syndrome 2, Breast and colorectal cancer | Cancer of multiple types (associated with ClinVar search term "Renal Cancer") |  |
| ***CHEK2*** | c.470T>C | Adrenocortical carcinoma, Gastrointestinal carcinoma, Hereditary cancer-predisposing syndrome, Prostate cancer, susceptibility to, Familial cancer of breast, Breast cancer, susceptibility to, Colon cancer, susceptibility to, Breast cancer, susceptibility to, Colorectal cancer, susceptibility to, Li-Fraumeni syndrome 2, Breast and colorectal cancer | Cancer of multiple types (associated with ClinVar search term "Renal Cancer") |  |
| ***CHEK2*** | c.1427C>T | CHEK2-Related Cancer Susceptibility, Colorectal cancer, Hereditary cancer-predisposing syndrome, Neoplasm of the breast, Breast and colorectal cancer, susceptibility to, Familial cancer of breast |  |  |
| ***CHEK2*** | c.1100DELC | Hereditary cancer, CHEK2-Related Cancer Susceptibility, Li-Fraumeni syndrome, Breast and colorectal cancer, susceptibility to, Familial cancer of breast, Breast cancer, susceptibility to, B Lymphoblastic Leukemia/Lymphoma, Not Otherwise Specified, Diffuse intrinsic pontine glioma, Ovarian Neoplasms, Leiomyosarcoma, Neoplasm of the breast, Astrocytoma, Li-Fraumeni syndrome 2, Malignant tumor of prostate, Osteosarcoma, Familial cancer of breast, Hereditary cancer-predisposing syndrome, Neoplasm of the breast |  | Thrombocytopenia, Colitis, Inflammation of the large intestine, Hematochezia |
| ***CHEK2*** | c.1100DELC | Hereditary cancer, CHEK2-Related Cancer Susceptibility, Li-Fraumeni syndrome, Breast and colorectal cancer, susceptibility to, Familial cancer of breast, Breast cancer, susceptibility to, B Lymphoblastic Leukemia/Lymphoma, Not Otherwise Specified, Diffuse intrinsic pontine glioma, Ovarian Neoplasms, Leiomyosarcoma, Neoplasm of the breast, Astrocytoma, Li-Fraumeni syndrome 2, Malignant tumor of prostate, Osteosarcoma, Familial cancer of breast, Hereditary cancer-predisposing syndrome, Neoplasm of the breast |  | Thrombocytopenia, Colitis, Inflammation of the large intestine, Hematochezia |
| ***CHEK2*** | c.1283C>T | Hereditary cancer-predisposing syndrome, Neoplasm of the breast |  |  |
| ***CHEK2*** | c.1100DELC | Breast cancer, susceptibility to, CHEK2-Related Cancer Susceptibility, Colorectal cancer |  |  |
| ***CHEK2*** | c.1100DELC | Li-Fraumeni syndrome 2, Malignant tumor of prostate, Osteosarcoma |  |  |
| ***CHEK2*** | c.349A>G | Familial cancer of breast | Hereditary cancer-predisposing syndrome (associated with ClinVar search term "Renal Cancer") |  |
| ***CHEK2*** | c.1283C>T | Neoplasm of the breast, Breast cancer, susceptibility to, CHEK2-Related Cancer Susceptibility, Colorectal cancer, Li-Fraumeni syndrome 2, Malignant tumor of prostate, Osteosarcoma, Familial cancer of breast, Hereditary breast and ovarian cancer syndrome, Breast and colorectal cancer, susceptibility to, Familial cancer of breast | Hereditary cancer-predisposing syndrome (associated with ClinVar search term "Renal Cancer") |  |
| ***CHEK2*** | c.1100DELC | Hereditary cancer, CHEK2-Related Cancer Susceptibility, Li-Fraumeni syndrome, Breast and colorectal cancer, susceptibility to, Familial cancer of breast, Breast cancer, susceptibility to, B Lymphoblastic Leukemia/Lymphoma, Not Otherwise Specified, Diffuse intrinsic pontine glioma, Ovarian Neoplasms, Leiomyosarcoma, Neoplasm of the breast, Astrocytoma, Li-Fraumeni syndrome 2, Malignant tumor of prostate, Osteosarcoma, Familial cancer of breast, Hereditary cancer-predisposing syndrome, Neoplasm of the breast |  | Thrombocytopenia, Colitis, Inflammation of the large intestine, Hematochezia |
| ***CHEK2*** | c.1100DELC | Hereditary cancer, CHEK2-Related Cancer Susceptibility, Li-Fraumeni syndrome, Breast and colorectal cancer, susceptibility to, Familial cancer of breast, Breast cancer, susceptibility to, B Lymphoblastic Leukemia/Lymphoma, Not Otherwise Specified, Diffuse intrinsic pontine glioma, Ovarian Neoplasms, Leiomyosarcoma, Neoplasm of the breast, Astrocytoma, Li-Fraumeni syndrome 2, Malignant tumor of prostate, Osteosarcoma, Familial cancer of breast, Hereditary cancer-predisposing syndrome, Neoplasm of the breast |  | Thrombocytopenia, Colitis, Inflammation of the large intestine, Hematochezia |
| ***CHEK2*** | c.1100DELC | Hereditary cancer, CHEK2-Related Cancer Susceptibility, Li-Fraumeni syndrome, Breast and colorectal cancer, susceptibility to, Familial cancer of breast, Breast cancer, susceptibility to, B Lymphoblastic Leukemia/Lymphoma, Not Otherwise Specified, Diffuse intrinsic pontine glioma, Ovarian Neoplasms, Leiomyosarcoma, Neoplasm of the breast, Astrocytoma, Li-Fraumeni syndrome 2, Malignant tumor of prostate, Osteosarcoma, Familial cancer of breast, Hereditary cancer-predisposing syndrome, Neoplasm of the breast |  | Thrombocytopenia, Colitis, Inflammation of the large intestine, Hematochezia |
| ***CHEK2*** | c.444+1G>A | CHEK2-Related Cancer Susceptibility, Li-Fraumeni syndrome 2, Malignant tumor of prostate, Familial cancer of breast, Osteosarcoma, Hereditary cancer-predisposing syndrome, not provided, Breast and colorectal cancer, susceptibility to, Familial cancer of breast, Breast cancer |  |  |
| ***CHEK2*** | c.1100DELC | Hereditary cancer, CHEK2-Related Cancer Susceptibility, not provided, Li-Fraumeni syndrome, Breast and colorectal cancer, susceptibility to, Familial cancer of breast, Breast cancer, susceptibility to, B Lymphoblastic Leukemia/Lymphoma, Not Otherwise Specified, Diffuse intrinsic pontine glioma, Ovarian Neoplasms, Leiomyosarcoma, Neoplasm of the breast, Astrocytoma, Li-Fraumeni syndrome 2, Malignant tumor of prostate, Osteosarcoma, Familial cancer of breast, Hereditary cancer-predisposing syndrome, Neoplasm of the breast |  | Thrombocytopenia, Colitis, Inflammation of the large intestine, Hematochezia |
| ***FH*** | c.1431_1433DUPAAA | Hereditary cancer-predisposing syndrome |  | Fumarase deficiency |
| ***FH*** | c.1431_1433DUPAAA | Hereditary cancer-predisposing syndrome |  | Fumarase deficiency |
| ***FLCN*** | c.619-1G>A | Hereditary cancer-predisposing syndrome | Multiple fibrofolliculomas |  |
| ***FLCN*** | c.1153DELC |  |  | not reported |
| ***FLCN*** | c.1429C>T | Carcinoma of colon, Hereditary cancer-predisposing syndrome | Renal cell carcinoma, nonpapillary, Multiple fibrofolliculomas | Pneumothorax, primary spontaneous, Chromosome 17, trisomy 17p11 2 |
| ***FLCN*** | c.619-1G>A | Hereditary cancer-predisposing syndrome | Multiple fibrofolliculomas |  |
| ***FLCN*** | c.1285DUPC | Hereditary cancer-predisposing syndrome | Multiple fibrofolliculomas | Pneumothorax, primary spontaneous |
| ***MET*** | c.3754T>C |  |  | not reported |
| ***MITF*** | c.952G>A | Cutaneous malignant melanoma 8 | Hereditary cancer-predisposing syndrome (associated with ClinVar search term "Renal Cancer") | Tietz syndrome, Waardenburg syndrome type 2A |
| ***MITF*** | c.952G>A | Cutaneous malignant melanoma 8 | Hereditary cancer-predisposing syndrome (associated with ClinVar search term "Renal Cancer") | Tietz syndrome, Waardenburg syndrome type 2A |
| ***MITF*** | c.952G>A | Cutaneous malignant melanoma 8 | Hereditary cancer-predisposing syndrome (associated with ClinVar search term "Renal Cancer") | Tietz syndrome, Waardenburg syndrome type 2A |
| ***MLH1*** | c.2252_2253DELAA |  |  | not reported |
| ***MRE11A*** | c.1090C>T |  |  | not reported |
| ***MRE11A*** | c.1090C>T |  |  | not reported |
| ***MRE11A*** | c.1090C>T |  |  | not reported |
| ***MSH2*** | c.1216C>T | Hereditary cancer-predisposing syndrome, Lynch syndrome I, Hereditary nonpolyposis colon cancer, Turcot syndrome, Muir-Torré syndrome, Lynch syndrome I, Lynch syndrome, Carcinoma of colon |  |  |
| ***MSH2*** | EX1_7inv |  |  | not reported |
| ***MSH6*** | c.1238G>A | Hereditary cancer-predisposing syndrome |  |  |
| ***MSH6*** | c.2731C>T | Turcot syndrome, Hereditary nonpolyposis colorectal cancer type 5, Endometrial carcinoma, Hereditary nonpolyposis colorectal cancer type 5, Hereditary nonpolyposis colon cancer, Hereditary cancer-predisposing syndrome, Lynch syndrome, Endometrial carcinoma |  |  |
| ***MSH6*** | c.3261DUPC |  |  | not reported |
| ***MUTYH*** | c.91DELG |  |  | not reported |
| ***MUTYH*** | c.536A>G |  |  | not reported |
| ***MUTYH*** | c.1187G>A |  |  | not reported |
| ***MUTYH*** | c.536A>G |  |  | not reported |
| ***MUTYH*** | c.1187G>A |  |  | not reported |
| ***MUTYH*** | c.1187G>A |  |  | not reported |
| ***MUTYH*** | c.963DELG |  |  | not reported |
| ***MUTYH*** | c.536A>G |  |  | not reported |
| ***MUTYH*** | c.358DELG |  |  | not reported |
| ***MUTYH*** | c.1187G>A |  |  | not reported |
| ***MUTYH*** | c.1187G>A |  |  | not reported |
| ***MUTYH*** | c.1214C>T |  |  | not reported |
| ***MUTYH*** | c.462+2T>G | Hereditary cancer-predisposing syndrome |  |  |
| ***MUTYH*** | c.1187G>A |  |  | not reported |
| ***MUTYH*** | c.1187G>A |  |  | not reported |
| ***NBN*** | c.2238DELC |  |  | not reported |
| ***NBN*** | c.657_661DELACAAA |  |  | not reported |
| ***NBN*** | c.657_661DELACAAA |  |  | not reported |
| ***NF1*** | 5'UTR_3'UTRdel |  |  | not reported |
| ***PALB2*** | c.2711G>A | Hereditary cancer-predisposing syndrome, Familial cancer of breast |  |  |
| ***PALB2*** | c.3113G>A | Hereditary cancer-predisposing syndrome, Familial cancer of breast, PALB2-Related Disorders, Hereditary breast and ovarian cancer syndrome, Breast cancer |  |  |
| ***PALB2*** | EX13_3'UTRdup |  |  | not reported |
| ***PALB2*** | c.758DUPT | Hereditary cancer-predisposing syndrome, Familial cancer of breast |  |  |
| ***PMS2*** | c.137G>T | Hereditary nonpolyposis colon cancer, Hereditary cancer-predisposing syndrome, Hereditary nonpolyposis colorectal cancer type 4, Turcot syndrome, Lynch syndrome, Hereditary nonpolyposis colorectal cancer type 4, Turcot syndrome, Pituitary carcinoma |  |  |
| ***PMS2*** | c.736_741DELCCCCCTINSTGTGTGTGAAG |  |  | not reported |
| ***PMS2*** | c.2155C>T | Hereditary cancer-predisposing syndrome |  |  |
| ***PMS2*** | c.137G>T | Hereditary nonpolyposis colon cancer, Hereditary cancer-predisposing syndrome, Hereditary nonpolyposis colorectal cancer type 4, Turcot syndrome, Lynch syndrome, Hereditary nonpolyposis colorectal cancer type 4, Turcot syndrome, Pituitary carcinoma |  |  |
| ***PMS2*** | c.1A>G | Hereditary nonpolyposis colon cancer, Hereditary cancer-predisposing syndrome, Lynch syndrome, Hereditary nonpolyposis colorectal cancer type 4, Lynch syndrome I |  |  |
| ***PTEN*** | c.49C>T | Hereditary cancer-predisposing syndrome, PTEN hamartoma tumor syndrome |  |  |
| ***PTEN*** | c.209+4_209+7DELAGTA | Ovarian Neoplasms, Cowden syndrome 1, Hereditary cancer-predisposing syndrome, PTEN hamartoma tumor syndrome |  |  |
| ***PTEN*** | c.48T>A | Hereditary cancer-predisposing syndrome, PTEN hamartoma tumor syndrome |  |  |
| ***PTEN*** | c.46DUPT |  |  | not reported |
| ***PTEN*** | in1_3'UTRdel |  |  | not reported |
| ***RAD51C*** | c.905-2_905-1DELAG | Breast-ovarian cancer, familial 3, Fanconi anemia, complementation group O, Hereditary cancer-predisposing syndrome |  |  |
| ***SDHA*** | c.1432_1432+1DELGG |  |  | not reported |
| ***SDHA*** | c.91C>T | Hereditary cancer-predisposing syndrome, Pilocytic astrocytoma | Paragangliomas 5, Carney triad | Mitochondrial complex II deficiency, Leigh syndrome, Dilated cardiomyopathy |
| ***SDHB*** | c.725G>A | Gastrointestinal stroma tumor, Gastrointestinal stroma tumor, Hereditary cancer-predisposing syndrome | Pheochromocytoma, Paragangliomas 4, Hereditary Paraganglioma-Pheochromocytoma Syndromes |  |
| ***SDHB*** | c.380T>G | Gastrointestinal stroma tumor, Gastrointestinal stroma tumor, Hereditary cancer-predisposing syndrome | Hereditary Paraganglioma-Pheochromocytoma Syndromes, Pheochromocytoma, Paragangliomas 4, Carney triad, Hereditary Paraganglioma-Pheochromocytoma Syndromes |  |
| ***TP53*** | c.827C>A | Hereditary cancer-predisposing syndrome |  |  |
| ***VHL*** | 5'UTR_3'UTRdel |  |  | not reported |
| ***VHL*** | c.266T>C | Hereditary cancer-predisposing syndrome | Von Hippel-Lindau syndrome | Erythrocytosis, familial, 2, |
| ***VHL*** | c.238A>G | Hereditary cancer-predisposing syndrome |  |  |
| ***VHL*** | c.232A>T | Hereditary cancer-predisposing syndrome | Von Hippel-Lindau syndrome |  |
| ***VHL*** | c.481C>T | Hereditary cancer-predisposing syndrome | Von Hippel-Lindau syndrome, Renal cell carcinoma, papillary, 1 | Erythrocytosis, familial, 2 |

**Supplemental Table 8. Odds Ratios of pathogenic gene variants for specific renal cancer histology**

| Gene | Chromo-phobe | Papillary renal | Clear cell | Wilms | Renal cell | Unknown | Other | Mixed Papillary | Mixed Chromo-phobe | Mixed  Oncocytoma |
| --- | --- | --- | --- | --- | --- | --- | --- | --- | --- | --- |
| *APC* | **-** | - | 1.24 | - | 2.02 | 0.94 | - | - | - | - |
| *ATM* | - | - | 3.13 | - | 1.15 | 0.23 | - | 10.22 | - | - |
| *BARD1* | - | - | 2.48 | - | - | 1.88 | - | - | - | - |
| *BRCA1* | - | - | 1.83 | 7.7 | - | 1.45 | - | - | - | - |
| *BRCA2* | - | 2.37 | 0.27 | - | 2.77 | 1.28 | - | - | - | - |
| *BRIP1* | - | - | - | - | 4.03 | 1.88 | - | - | - | - |
| *CHEK2* | 2.43 | - | 1.14 | 2.49 | 0.75 | 1.1 | - | - | - | - |
| *MSH6* | - | - | 0.82 | - | 4.05 | 0.63 | - | - | - | - |
| *MUTYH* | - | - | 1.24 | 22.49 | - | - | - | - | - | 184.73 |
| *NBN* | 10.2 | - | - | - | - | 3.77 | - | - | - | - |
| *PALB2* | - | - | 2.48 | - | - | 1.89 | - | - | - | - |
| *PMS2* | - | - | 0.49 | - | 0.8 | 3.79 | - | - | - | - |
| *BAP1* | - | - | 1.19 | - | - | 4.16 | - | - | - | - |
| *FLCN* | 4.29 | - | - | - | 1.03 | 3.48 | - | - | - | - |
| *MITF* | - | - | 1.07 | - | - | 1.15 | - | - | 102.37 | - |
| *PTEN* | 5.11 | - | 0.62 | - | - | 2.84 | - | - | - | - |
| *SDHB* | - | - | 2.14 | - | - | - | - | - | - | 275.91 |
| *VHL* | - | - | 0.53 | - | 2.76 | 0.57 | 10.19 | - | - | - |
| Odds ratio > 1 = higher than expected due to chance alone, Odds ratio < 1 = lower than expected due to chance alone; - = calculations unable to be completed due to a value of zero. Blue font – RC-specific genes, black font- other cancer genes, and bold black- DDR genes. Genes with Odds ratio of 0 for all categories are not listed in the table. | | | | | | | | | | |
